# East-African savanna dynamics: from a knowledge-based model to the possible futures of a social-ecological system

**DOI:** 10.1101/2021.04.05.438440

**Authors:** Maximilien Cosme, Christelle Hély, Franck Pommereau, Paolo Pasquariello, Christel Tiberi, Anna Treydte, Cédric Gaucherel

**Author notes:** Corresponding author: Maximilien Cosme. UMR AMAP, CIRAD - TA A51/PS1, 34398 Montpellier cedex 5 - France.

## Abstract

Sub-Saharan savanna ecosystems are undergoing transitions such as bush encroachment, desertification or agricultural expansion. Such shifts and persistence of land cover are increasingly well understood, especially bush encroachment which is of major concern in pastoral systems. Although dominant factors can explain such transformations, they often result from intertwined causes in which human activities play a significant role. Therefore, in this latter case, these issues may require integrated solutions, involving many interacting components. Ecosystem modelling has proved appropriate to support decision-makers in such complex situations. However, ecosystem models often require lots of quantitative information for estimating parameters and the precise functional form of interactions is often unknown. Alternatively, in rangeland management, States-and-Transitions Models (STMs) have been developed to organize knowledge about system transitions and to help decision-makers. However, these conceptual diagrams often lack mathematical analyzing tools, which strongly constrains their complexity. In this paper, we introduce the Ecological Discrete-Event Network (EDEN) modelling approach for representing the qualitative dynamics of an East-African savanna as a set of discrete states and transitions generated from empirical rules. These rules are derived from local knowledge, field observations and scientific literature. In contrast with STMs, EDEN generates automatically every possible states and transitions, thus enabling the prediction of novel ecosystem structures. Our results show that the savanna is potentially resilient to the disturbances considered. Moreover, the model highlights all transitions between vegetation types and socio-economic profiles under various climatic scenarios. The model also suggests that wildlife diversity may increase socio-economic resistance to seasonal drought. Tree-grass coexistence and agropastoralism have the widest ranges of conditions of existence of all vegetation types and socio-economic profiles, respectively. As this is a preliminary use of EDEN for applied purpose, analysis tools should be improved to enable finer investigation of desirable trajectories. By translating local knowledge into ecosystem dynamics, the EDEN approach seems promising to build a new bridge between managers and modellers.

## Introduction

Among the great variety of East African landscapes, savannas are critical ecosystems for human populations. Indeed, throughout East Africa, human societies benefit from these highly productive systems (Ryan et al., 2016) and while climate and wildlife are known as an important driver of savanna dynamics (Ellis and Swift, 1988), it is also widely acknowledged that human activities such as agriculture, pastoralism and tourism contribute to land conversion into reserves, cities, croplands or rangelands (Little, 1996; Marchant et al., 2018; Pratt et al., 1977). These latter changes strongly affect biodiversity and natural resources distribution, but may also undermine the persistence of human activities (e.g. pastoralism; Liao and Clark, 2018) themselves. Therefore, anticipating the far-reaching consequences of such changes in East African social-ecological systems is key for improving societies’ adaptive capacity.

Ecological models are used for predicting such consequences to various drivers, such as variations in rainfall regime (Hély et al., 2006) or herbivore populations increase (Hein, 2006; Sinclair et al., 2015). So far, many models consider a few drivers simultaneously and rarely take qualitative expert knowledge (Bashari et al., 2008) or historical anecdotes (Archer, 1989) into account. The predictive power of such quantitative models is thus highly limited by the acquisition cost of quantitative data (Gaucherel et al., 2011; Kauffman, 1969; Levins, 1966; Thomas, 1991). Yet, a deep qualitative knowledge on social-ecological system functioning can be derived from local people such as pastoral communities, land managers and farmers (Bélisle et al., 2018). Qualitative models using such expert-knowledge display interesting features for ecological questions (Dambacher et al., 2003; Gaucherel et al., 2017; Salles and Bredeweg, 2006; Starfield, 1990; Westoby et al., 1989; Zhao et al., 2011) and can be complementary to quantitative models (Rykiel, 1989). However, qualitative models imply sacrificing system behaviors’ precision to gain realism and generality (Levins, 1966). Realism can be gained through a more integrated model (i.e. including various scientific and non-scientific sources of knowledge) (Gaucherel et al., 2017), while the reduction of parametric sensitivity increases model generality (Hosack et al., 2008; Puccia and Levins, 1991; Saadatpour and Albert, 2016). In addition, a qualitative approach enables a more comprehensive assessment of the *possible* system responses to any stress or disturbance (Gaucherel and Pommereau, 2019; Plant et al., 1999), which could prove useful for addressing management issues (Bestelmeyer et al., 2004; Cordier et al., 2014; Suding and Hobbs, 2009).

Tropical savannas vegetation and their dynamics have been described qualitatively worldwide (Bodini and Clerici, 2016; Hirota et al., 2011; Sinclair, 2008). They usually display three distinct qualitative states, namely grassland, savanna and woodland (Hirota et al., 2011; Noy-Meir, 1975; Scholes and Archer, 1997; Walker et al., 1981), with bare soil as the unvegetated state, and only rarely display intermediate states (Sala and Maestre, 2014). Still today, the mechanisms lying behind their conditions of existence and stability are still debated (Baudena et al., 2015; Staver and Levin, 2012; Touboul et al., 2018). Moreover, some quantities such as soil water content, grass biomass or herbivores density vary dramatically between seasons and may support a qualitative description (e.g. low/high). These characteristics are encouraging for complementing the quantitative approaches by adopting a qualitative description of the dynamics of tropical savannas.

In rangeland ecology, a kind of qualitative model called States-and-Transition Models (STMs) have been proposed as a support for decision-makers in rangeland management and to account for irreversible transitions in vegetation (Briske et al., 2005; Westoby et al., 1989), as for instance in Ethiopian savannas (Liao and Clark, 2018). STMs are conceptual box-and-arrow diagrams representing the system dynamics as discrete states and irreversible transitions, where each state is defined by the reversible transitions between community phases (Stringham et al., 2003). States and transitions composing the STMs are derived from empirical observations, making these models highly relevant for management actions in specific contexts. However, this strong reliance on empirical observations limits their usefulness if new ecosystems are emerging (the so-called novel ecosystems; Morse et al., 2014). A major goal would thus be, in Walker and Westoby (2020) terms, to “build a constructive ecology where novel ecosystems can be imagined, but in a disciplined way, restricted by well-founded knowledge about processes and transitions”. Combining local knowledge and mathematical analysis would thus allow managers and modellers to anticipate new transitions, new ecosystem structures and the far-reaching consequences of social and ecological changes in the system (Peinetti et al., 2019; Plant et al., 1999).

Among qualitative models, discrete-event models (Cassandras and Lafortune, 2008) have shown their ability to support management decision-making (Cordier et al., 2014), to handle complexity rigorously and fruitfully (Chabrier and Fages, 2003; Gaucherel et al., 2017; Gronewold and Sonnenschein, 1998; Thomas, 1991; Zhao et al., 2011) and to provide conceptual advances in systems dynamics (Melliti et al., 2015; Thomas, 1981). These models represent the system dynamics as discrete event-driven changes (i.e. at possibly irregular time steps) and many methods enable tracking the complex cascading effects of transitions (Monteiro et al., 2008). In addition, like STMs, they often provide a graphical and intuitive representation of social-ecological systems dynamics, thus improving communication of complex phenomena towards the non-scientific public such as stakeholders and students (Bestelmeyer et al., 2009; Cassandras and Lafortune, 2008).

In this study, we introduce the *Ecological Discrete-Event Network modelling approach* (EDEN for short) (Gaucherel and Pommereau, 2019) to assess the potential resilience (Grimm and Wissel, 1997) as well as vegetation and socio-economic qualitative dynamics of a generic East-African savanna ecosystem. The computed outputs of EDEN share the same structure as the aforementioned STMs (Bestelmeyer et al., 2009; Westoby et al., 1989), although EDEN computes states and transitions automatically from ecosystem variables and their interactions instead of defining them based on empirical observations and expert knowledge (Plant et al., 1999). Based on field surveys in Northern Tanzania, expert knowledge (from Istituto Oikos, Nelson Mandela African Institution of Science and Technology) and scientific literature (e.g. Bobe, 2006; Ogutu et al., 2008; Sankaran et al., 2008), we built this qualitative model to assess: (i) the potential resilience of the system when it faces various perturbations alone or combined, (ii) the range of social-ecological conditions of key ecosystem variables and profiles (i.e. combinations of variables) and (iii) the transitions between these profiles. Here, we will propose the concept of “potential resilience”. It is a qualitative property indicating that the initial state remains reachable after any transition (independently of recovery time and transitions probabilities), although several transient drastic shifts in vegetation structure and/or socio-economy may occur. Although resilience may be influenced by the spatial structure (Allen et al., 2016), the model presented here is spatially implicit, assuming that it still enables testing the following hypotheses.

First, in the long term and given the considered variables and transitions of such a social-ecological system, we hypothesized that (H1) the savanna system is potentially resilient in the face of most disturbances (Gil-Romera et al., 2010). Secondly, tree-grass coexistence and agro-pastoral societies are widespread features of past and current East-African landscapes (Lankester and Davis, 2016). Therefore, (H2) we expect tree-grass coexistence and agro-pastoralism to exist in a broader range of conditions than others vegetation types and socio-economic profiles, respectively (House et al., 2003; Nelson et al., 2009).

## Material and methods

### Field surveys

Field surveys were conducted in 2018 in Arusha region, Northern Tanzania. In this region, annual precipitation is bimodal, with a long and a short rainy season (from March to May and November to December, respectively). Mean annual precipitation is about 700 mm (Gereta et al., 2004). Two sites were surveyed, the Gelai plains and Meru savanna, in order to get a broad view of northern-Tanzanian savannas, ranging from open to wooded savannas. Gelai plains (2°47’52.1”S, 36°06’03.1”E) are grassy plains located between the mount Kitumbeine and Lake Natron, mostly used for tourism and pastoralism. Meru savannas are located north to the mount Meru volcano (3°09’34.5”S, 36°46’53.2”E). They consist of a mosaic of grasslands, savannas and woodlands, where agro-pastoral activities are practiced (Istituto Oikos, 2011). We did not find any fire scars on tree trunks nor charcoals attesting recent fire on any site. More information about study area is available in Supplementary material (see Appendix S1 in Supplementary material). These surveys served as an additional source of information to the scientific literature in order to build a generic picture of East-African rangelands.

### Discrete-event and rule-based modelling

Most models in ecology represent time as regular intervals between events. This is the case with continuous-time models (differential equations or continuous-time Markov chains) or discrete time models (discrete Markov Chains, difference equations). In contrast, discrete-events models describe system dynamics in terms of *events*, i.e. abrupt physical changes possibly occurring at irregular intervals (Cassandras and Lafortune, 2008). These models, and most notably Asynchronous Boolean Networks (Thomas, 1991), have been used for a long time in systems biology to model regulation networks (Chabrier and Fages, 2003; Kauffman, 1969; Thomas, 1991, 1973), but only recently in ecology (Baldan et al., 2015; Campbell et al., 2015; Gaucherel and Pommereau, 2019; but see Gronewold and Sonnenschein, 1998). In the EDEN approach, the values of ecological interaction parameters are not required (i.e. the modelled dynamics are parameter-insensitive). This enables the derivation of robust predictions by the sole use of qualitative data available from the various stakeholders and in the literature. Moreover, the lower computational cost in qualitative models facilitates the exploration of the full range of possible behaviors. Finally, as the model only consists of discrete states and transitions, the chronology of successive events is preserved. Variables may represent biotic, abiotic or anthropogenic entities and are *Boolean* (i.e. binary). A system *state* is described by its present variables within brackets as {variable1, variable2}. Therefore, a state in EDEN should not be confused with states in STMs, which are defined by a set of reversible community transitions (Stringham et al., 2003).

Building on an historical and fruitful use of Boolean Networks in systems biology (Kauffman, 1969; Thomas, 1973), the Boolean modelling of ecological systems has already proved useful in theoretical studies, especially when the precise functional form and parameters of ecological interactions are unknown (Joo et al., 2018). For instance, Robeva and Murrugarra (2016) showed that a simple synchronous Boolean network can reproduce the bistable behavior observed in traditional models of budworm-forest dynamics (Holling, 1973; Ludwig et al., 1978). Boolean models of mutualistic networks also yielded insights about community assembly (Campbell et al., 2011), response to extinctions (Campbell et al., 2012) and invasibility (Campbell et al., 2015). More recently, discrete-event Boolean models (Gaucherel et al., 2020; Mao et al., 2021) have been used for ecosystem services assessment, further expanding their use in ecology.

But what could justify the use of a Boolean abstraction in an ecological context? The answer lies in non-linear ecological phenomena, which appear ubiquitous in biology ecology (D’Amario et al., 2019; Thomas, 1991). In such phenomena, a variable may exhibit a different response to driver whether the latter is above or below a given threshold. Below the threshold, where the variable is not or slightly responding, the driver can be considered “functionally absent” and valued by a “-”. Conversely, above the threshold, the driver is considered “functionally present” (noted “+”) and causes a strong, qualitative change in variable response. As a simple ecological illustration, consider seeds germination (the variable) triggered by soil moisture (the driver). Above a given - and often unknown (D’Amario et al., 2019) - threshold of soil moisture, the latter has a qualitative effect on seed germination (Hillel, 1972). In our model, we assume that such a threshold exists for each interaction represented in the model. In summary, by focusing on variables presence and absence, we focus on the most abrupt changes in the ecological system structure and ignore “below-threshold” quantitative variations. Initially, the modeller defines a set of variables, their initial state and the set of transitions rules, as it has been done previously in other models (Salles and Bredeweg, 2006; Starfield, 1990). The method applied here follows the work of Gaucherel and Pommereau (2019).

The EDEN approach includes a formalism based on transition rules comprised of a *condition* and a *realization* (Table 1). A condition is a subset of variables allowing the execution of a rule. The realization of this rule then changes variables values, thus creating a new ecosystem state. This process is symbolized by an arrow (→ or ≫) (Table 1). Given a transition from a state (or set of states) A to a state (or set of states) B, we say that A is a predecessor of B, and B a successor of A. A rule is executed every time its condition is satisfied in a given ecosystem state. Moreover, priority rules (called constraints) can also be used to model fast processes. More details on transitions, rules, constraints and their firing conditions can be found in Appendix S2.

**Table. 1.** Rule set of the “Grasses-Trees-Livestock” toy-model. The variable Gr (Grasses) refers to any grass species in the system, Tr (Trees) to any woody plant and Lv (Livestock) to a grass-eating livestock species, e.g. cattle.

| Rule | Condition | Realization | Rule description |
| --- | --- | --- | --- |
| **R1** | Gr+ | → Lv+ | Grasses attract livestock. |
| **R2** | Gr+ | → Tr+ | Grasses promote trees growth. |
| **R3** | Lv+, Gr+ | → Tr - | Livestock may control trees if there is enough grass. |
| **R4** | Tr + | → Gr-, Lv- | Trees may outcompete grasses and thus exclude livestock. |
| **R5** | Tr - | → Gr+ | Grasses may reestablish if woodland is removed. |

Such a rule-based formalism can be found in various ecological models, including cellular automata, multi-agent systems or fuzzy logic. Generally, in these models, all executable rules in one state are executed simultaneously. Thus, all variables are updated at each time step or event. This property is called the *synchronous update mode*. This represents a strong assumption about ecological parameters as it implies that all variables change at the same speed (Garg et al., 2008), which strongly affects the possible dynamics, missing some event sequences (trajectories) and creating spurious ones. Alternatively, in EDEN, the system state is updated *asynchronously*: only one rule is executed at a time, which allows relaxing most assumptions on parameter values, and thus the possible trajectories (Garg et al., 2008). Asynchrony thus generates all the possible alternative trajectories in ecosystem dynamics. Therefore, the EDEN approach is called *possibilistic*. Possibilism is based on the idea that rare (yet possible) events should also be considered in ecosystem analysis, as they may have major consequences (e.g. irreversible shift in ecosystem structure) and would thus be highly relevant, e.g. in a management situation (Clarke, 2008). Possibilism is thus a non-probabilistic perspective on non-determinism (Clarke, 2008; Kristensen et al., 2019).

The EDEN approach is thus based on a qualitative, asynchronous and possibilistic model. The model output, which results from the successive firing of each possible transition until no more can be fired, is the state space (i.e. *ecosystem dynamics*). This state space is represented by a directed graph called the *State Transition Graph (STG)* in which nodes and edges represent system states and transitions, respectively (Fig. 1a). Details on STG computation can be found in (Gaucherel and Pommereau, 2019).

**Fig. 1.**
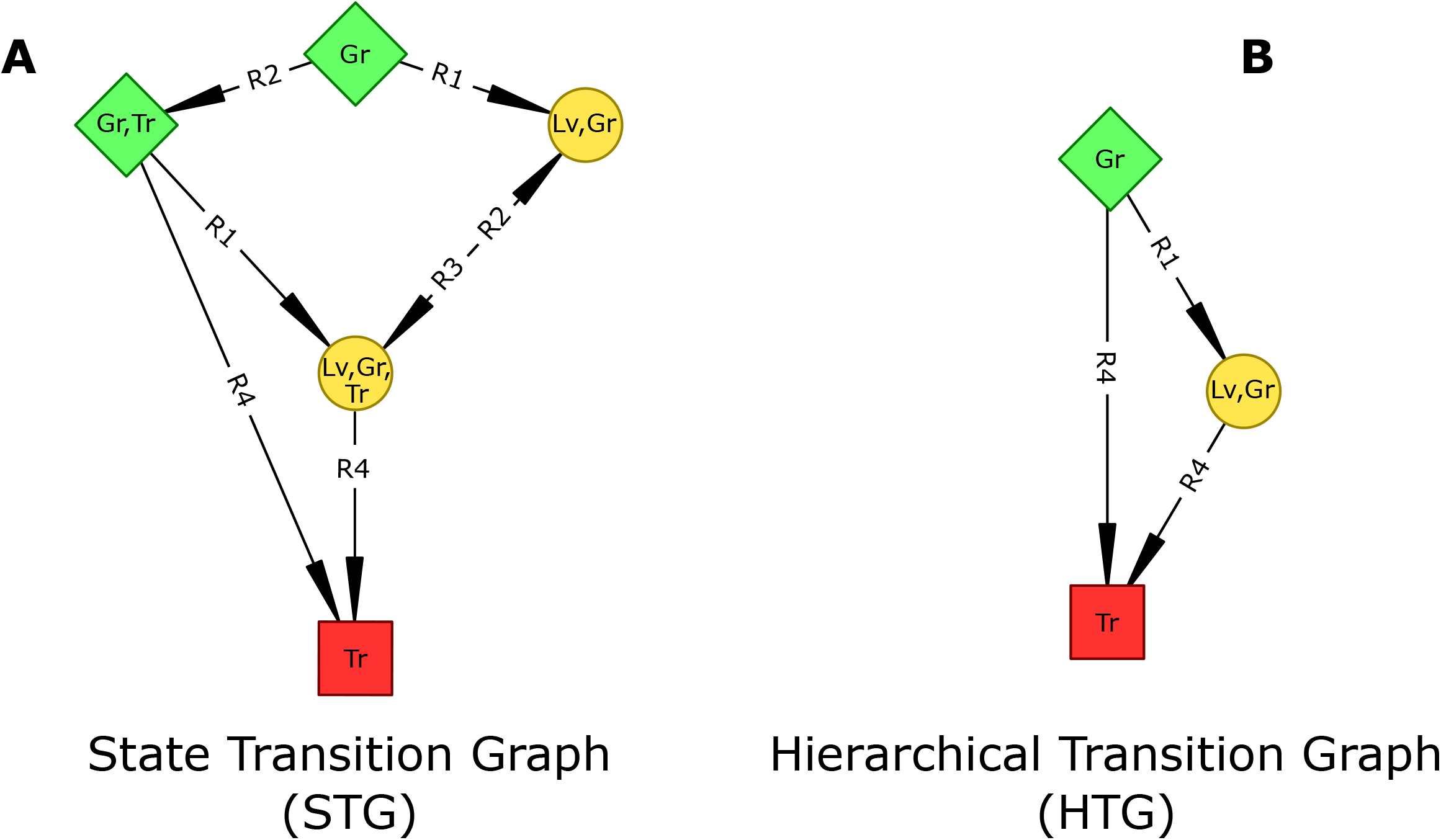
Transition graphs of the “Grasses-Trees-Livestock” toy-model. A: State Transition Graph (STG) depicting every reachable states and transitions from the initial state {Gr}. *B*: Hierarchical Transition Graph (HTG) depicting basins (green lozenge-shaped node), Strongly Connected Components or SCCs (yellow round-shaped node) and stable states (red square-shaped node). Note that the green basin (resp. yellow SCC) in the HTG results from the merging of the green states (resp. yellow states) in the STG. Rules labels and description are detailed in Table 1. Gr: Grasses; Tr: Trees; Lv: Livestock.

In the STG, we defined five topological structures and the corresponding ecosystem dynamics (Chaouiya et al., 2008; Naldi et al., 2011): (i) a Strongly Connected Component (SCC, depicted as a round node) represents a subset of states in which each state is reachable from any other state in this subset (Fig. 1a). Hence, by definition, every ecosystem change in a SCC is *reversible*, either directly or using a roundabout path. This concept is close to that of “state” in STMs (Westoby et al., 1989) or that of resilience (Grimm and Wissel, 1997) and will be used as such. A SCC can be transient or terminal (i.e. an attractor); (ii) an *attractor* is a SCC the system cannot exit. For instance, in systems biology, attractors may represent differentiated cell phenotypes (Bonzanni et al., 2014) ; (iii) a *transient SCC* is a non-terminal SCC (i.e. it is possible to exit it). The SCC containing the initial state (called initial SCC) and 1-state SCCs (called trivial SCCs) are subclasses of transient SCCs; (iv) a *stable state* (depicted as a square) is an individual state from which the system cannot get out. It is thus a special kind of attractor; (v) a *basin* (depicted as a lozenge) and is a set of states that can reach the same SCCs or stable states. They represent transient dynamics because they necessarily lead to a stable or oscillatory dynamics. As a STG can be very large (up to thousands of states), it can summarized by a Hierarchical Transition Graph (HTG) (Abou-Jaoudé et al., 2016; for formal definitions, see Bérenguier et al., 2013). Informally, a HTG is an acyclic graph whose nodes represent topological structures (i.e. SCCs, basins and stable states), with transitions indicating the existence of a path between two topological structures (Fig. 1b).

Equipped with these definitions, we can now precisely interpret the Fig. 1: The State Transition Graph (STG) represents all *possibly* reachable states from the initial state {Gr} (Fig. 1a). Yellow round-states are mutually reachable, and thus form a *Strongly Connected Component (SCC)*. The red square-state is a single state, has no successor and thus form a *stable state*. Green lozenge-states can reach the same SCC and stable state and thus form a *basin*. The Hierarchical Transition Graph (HTG) summarizes the STG by merging states belonging to the same basin or SCC (Fig. 1b), thus highlighting irreversible transitions in ecosystem dynamics. Note that the HTG does not indicate which state leads to the next topological structure (e.g. which state in the SCC leads to the stable state, Fig. 1), but only the variables remaining constant in the structure (Fig. 1b). The STG thus provides additional information about dynamics, and especially about reversible transitions. For instance, the periodic control of tree growth through livestock grazing (yellow states, Fig. 1a). Transitions can be labelled with rule number in order to provide a process-explicit view of system dynamics.

### The savanna model description

We considered a tropical savanna social-ecological system (Fig. S1). We built a reference model including 22 biotic, abiotic and socio-economic variables (Table 2), with 66 rules and 18 constraints representing system transitions (Table S1). The model was initially built based on field surveys and local knowledge. Local knowledge was acquired through interviews of various stakeholders (NGOs, biodiversity conservation scientists and farmers) and direct observation through field surveys. This knowledge was complemented with scientific literature about semi-arid vegetation and water dynamics, farming, the role of herbivory, pastoralism in East-Africa, human activities regulation and livestock-wildlife interactions. Disturbances (climatic, biological, ecological or anthropogenic) or management actions were considered intrinsic to the ecosystem (Rykiel, 1985) and, thus, represented as rules. We did not include fire in the model, as surveyed sites did not exhibit any recent sign of fire (i.e. no fire scar or black ash on tree trunks, nor burned grass clumps). In addition, satellite products such as NASA’s burned area product from the MODIS sensors clearly show that the region studied in northern Tanzania, and the region as a whole between Lake Victoria and the Indian Ocean is among the least burned areas in Africa despite the presence of vegetation. The initial state corresponds to the current state of the Meru site (Table 2).

**Table. 2.** Set of ecosystem variables. Reference: reference model version; M1: “Permanent dry season” model version; M2: “Permanent rainy season” model version. “+” or “-” indicate whether the variable is initially present or absent in the corresponding model version, respectively.

| Variable | Symbol | Description | Reference | M1 | M2 | M3 | M4 |
| --- | --- | --- | --- | --- | --- | --- | --- |
| Rainfall | **Rf** | Seasonal rainfall | + | - | + | + | + |
| Soil | **Sl** | Organo-mineral part of the ecosystem. | + | + | + | + | + |
| Temporary water | **Tw** | Rainfed or managers-fed water reservoirs | + | + | + | + | + |
| Aquifers | **Aq** | Deep water, higher residence time than temporary water. | + | + | + | + | + |
| Carnivores | **Ca** | Lions, hyaenas, cheetah... | + | + | + | + | + |
| Grazers | **Gz** | Wildebeests, zebras, antelopes... | + | + | + | + | + |
| Elephants | **El** | Elephant populations and mixed-feeders (e.g. impala) | + | + | + | + | - |
| Browsers | **Bw** | Giraffe, kudu, dik-dik... | + | + | + | + | - |
| Bovines | **Bv** | All bovine breeds. | + | + | + | - | + |
| Goats | **Go** | All goat breeds. | + | + | + | + | - |
| Sheeps | **Sh** | All sheep breeds. | + | + | + | - | + |
| Grasses | **Gr** | Herbaceous plants. | + | + | + | + | + |
| Trees | **Tr** | Woody plants, including trees, bushes and shrubs. | + | + | + | + | + |
| Crops | **Cr** | All crops. | + | + | + | + | + |
| Agriculturalists | **Ag** | Farmers sedentary population. | + | + | + | + | + |
| Settlements | **St** | Permanent houses or villages (Maasai bomas, small permanent structures). | + | + | + | + | + |
| Pastoralists | **Pa** | Nomadic or semi-nomadic populations living from crops and livestock. | + | + | + | + | + |
| Land managers | **Lm** | Any extra local authority that may act for biodiversity conservation and the functioning of human activities. | + | + | + | + | + |
| Tourists | **To** | External people coming to observe wildlife and Maasai culture. | + | + | + | + | + |
| Browsers diseases | **Bd** | Any pathogen affecting sheeps, goats and wild browsers. | - | - | - | - | - |
| Grazers diseases | **Gd** | Any pathogen affecting bovines and wild grazers (e.g. rinderpest). | - | - | - | - | - |
| Pests | **Ps** | Insects, crops parasites or fungi. | - | - | - | - | - |

### Analyses

First, we computed the mean ecosystem state, i.e. the frequency of each variable or group of variables over the states of the STG. We interpreted it as their range of social-ecological conditions of existence. In a second step, we focused on ecological and economic state properties that may appear relevant for managers. For that purpose, we defined a partition (Diop et al., 2019) of vegetation types (Scholes and Archer, 1997) and socio-economic profiles. A partition defines mutually exclusive groups of states including some present/absent variables. The partition of vegetation types includes four elements: bare soil (defined by Tr- and Gr-, i.e. no trees and no grasses), grassland (Tr- and Gr+), savanna (Tr+ and Gr+) and woodland (Tr+ and Gr-) (see Table 2 for variables labels). On the other hand, the partition of socio-economic profiles combines human activities (i.e. agriculture, noted A and defined by Ag+), pastoralism, P: Pa+; and tourism, T: To+) in eight elements: APT, AP, AT, PT, A, P, T and NH (No Humans) (Table 2). Note that human activities are defined by the corresponding human groups (agriculturalists, pastoralists, tourists), and not by activities per se.

A STG often includes thousands of states (Gaucherel et al., 2020; Mao et al., 2021), rendering visual analysis impracticable. Therefore, in a third step, we used the above-mentioned partitions to generate *summary graphs* (Diop et al., 2019; Monteiro et al., 2014). A summary graph summarizes specific aspects of the dynamics (e.g. vegetation dynamics) by focusing on the corresponding variables of interest (e.g. grasses and trees). It aggregates into one node all states belonging to the same group in the partition (e.g. all grassland states) (see Diop et al., 2020 for procedure) and highlights transitions between groups of the partition. Here, summary graphs of vegetation types and socio-economic activities will be called *vegetation summary graph* and *socio-economic summary graph*, respectively. This allowed computing the range of social-ecological conditions of existence for each of them by computing their respective size in comparison with the whole STG.

However, one could not only be interested in the existence of a group of states, such as grassland, but also in its potential persistence (e.g. could grassland persist or not?). Therefore, in some cases, we used *SCC-summary graphs*. Informally, this graph is constructed by (1) partitioning states (as explained earlier) according to some variables and then (2) merging states forming a SCC. The resulting SCC-summary graph is analog to States-and-Transitions Models used in rangeland management (Stringham et al., 2003; Westoby et al., 1989). Each SCC corresponds to a state (as defined in STMs, Stringham et al., 2003), although in our case SCCs are not restricted to vegetation states.

In a fourth step, we focused on SCCs averaged dynamics. Indeed, some variables in a SCC may be present in a few states, while others may be more frequent. To do so, we performed a Principal Components Analysis (PCA) based on variables’ frequency of occurrence in SCCs. Therefore, the frequency of every variable in a SCC will be called the *SCC profile*. The PCA will thus discriminate SCCs based on their profile. Finally, as a sensitivity analysis, we selectively removed key variables or rules to build four other model versions (Table 2; see two more versions in S2).

### Model versions

Seasonality is known to influence woody cover and vegetation transitions (Guttal and Jayaprakash, 2007; Hély et al., 2006). Hence, we removed the rule “Rf- → Rf+” (i.e. rainy-to-dry season shift) (model version n°1: M1, Table S4) and the rule “Rf+ → Rf-” (M2) from the reference model in order to assess the impacts of seasonality scenarios on savanna dynamics. As the delays in the onset of rainy season are of particular concern in tropical regions, the M1 scenario simulates a long-lasting dry season. It aims to assess the ultimate consequences of an arbitrarily long drought (i.e. onset of rainy season is maximally delayed) and identify the successive transformations of the social-ecological network induced by water stress. On the other hand, as increases in rainfall amount are expected in some tropical regions, M2 simulates a long lasting rainy season. Next, we removed grazers (wild grazers and cattle, M3) and browsers (wild browsers, elephants and goats, M4) independently (Table S4 and 5). Rules and/or variables were modified each model version to preserve ecological coherence (see Table S4 for details).

We hypothesized that a maximal delay in rainy season onset would lead to desertification (i.e. a widespread environmental degradation that reduces productivity of dryland ecosystems by, among other causes, reducing plant cover and soil loss; Whitford and Duval, 2020). On the other hand, a maximal delay in dry season onset would lead to a woodland development (Sankaran et al., 2005). Regarding the suppression of herbivory (M3, M4), as it is a well-known driver of vegetation changes at the community-level, potentially inducing large-scale vegetation shifts (Baumgartner et al., 2015; de Boer et al., 2015; Devine et al., 2017; Treydte et al., 2013), we expected vegetation transitions to be modified. Long-term anthropogenic effects on savanna dynamics are still debated (Sinclair et al., 2008). Hence, we removed all anthropogenic variables (here pastoralists, livestock, agriculturalists, crops, land managers, tourists and settlements) (Table S4, M5) and pastoral variables (Table S4, M6). Besides the removed variables from each version, all had the same initial state (Table 2).

## Results

### Reference model version (including all rules and variables)

The STG of the reference model showed 24,444 states and a unique SCC (Fig. 2a, see Table 2 for description of the initial state). Vegetation and socio-economic summary graphs highlighted specific aspects of its dynamics.

**Fig. 2.**
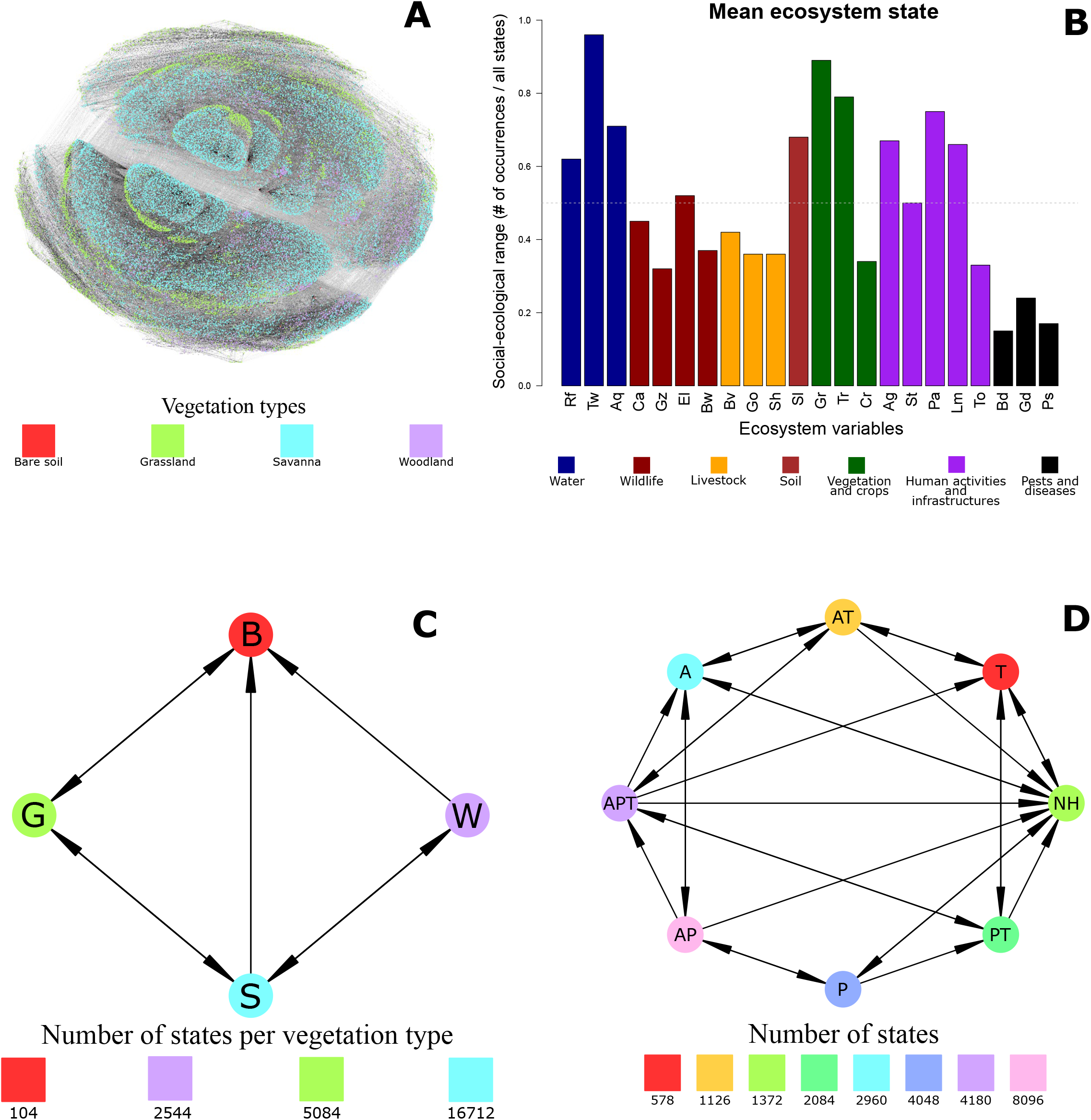
Reference model version. A: STG; B: “mean ecosystem state”, where each bar corresponds to the proportion of states in which the corresponding variable is present, i.e. the range of conditions of each variable. Variables descriptions are detailed in Table 2; C: Vegetation summary graph of Fig. 2a. Each node in this graph corresponds to all the system states of the corresponding vegetation type (B: bare soil; G: grassland; S: savanna; W: woodland); D: Socio-economic summary graph of Fig. 2a, based on socio-economic profiles. Each node corresponds to all the system states of the corresponding socio-economic profile (A: agriculture; P: pastoralism; T: tourism; NH: no humans; each letter combination correspond to a different socio-economic profile).

#### Vegetation

Grasses and trees showed almost the same range (Fig. 2b). Savanna exhibited the largest range (Fig. 2c). In the vegetation summary graph (Fig. 2c), bare soil appeared to be the only vegetation type reachable by any other type. Rules responsible for these vegetation transitions in Fig. 2c are detailed in Table S2.

#### Human activities and socio-economic profiles

Pastoralism showed the largest range (of social-ecological conditions of existence) and tourism the smallest one (Fig. 2b). The socio-economic summary graph (Fig. 2d) showed that the absence of humans (NH) is directly reachable by any socio-economic profile. Moreover, only one activity can be gained at a time, while several can be lost simultaneously.

Among socio-economic profiles, the agriculture-pastoralism combination had the largest range in all vegetation types except woodland and bare soil (where agriculture is dominant) (Table S7), while tourism still had the smallest range (2%). Rules responsible for these socio-economic transitions in Fig. 2d are detailed in Table S3. Note that the social-ecological range of a vegetation type or human activity is not a dynamical property, nor does it correspond to a probability of occurrence over time. It simply corresponds to the proportion of states displaying this vegetation type or human activity.

### “Permanent drought” version (M1)

Removing the rainy season rule (Rf- → Rf+) induced irreversible transitions, as indicated by the absence of ascending arcs in the HTG (Fig. 3 and S3). The system displays four transient SCCs and converges towards a stable state. From each SCCs, firing R6 or R8 (i.e. aquifers drying out) pushes the system into the stable state. Farmers’ migration (R42) may also play a similar role.

**Fig. 3.**
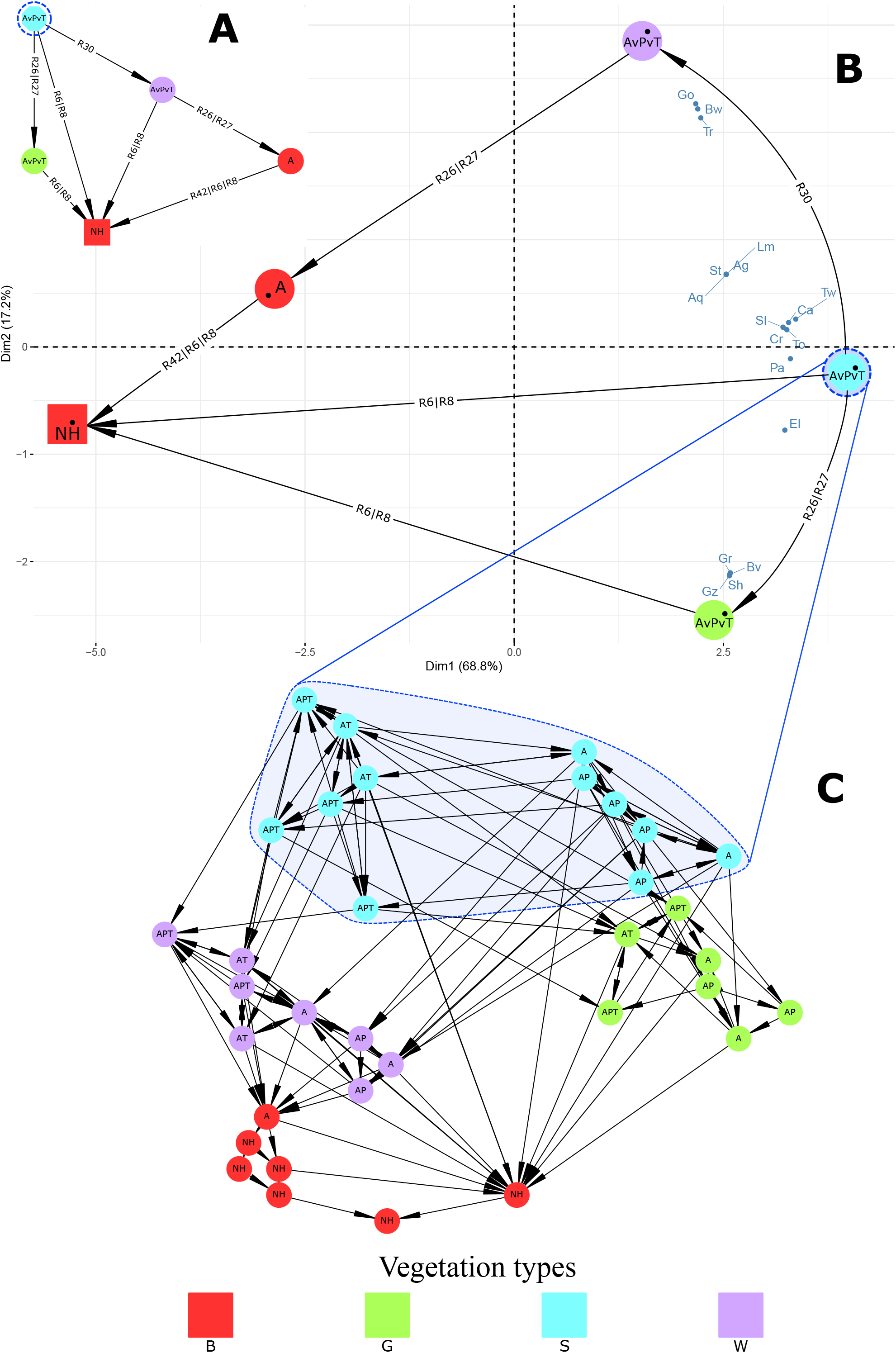
Sustained dry conditions (M1 model version). *A*: HTG, where each node is a topological structure (i.e. a SCC, basin or stable state). The graph is chronologically oriented from top to bottom. *B*: PCA projection of each SCC based on their composition; C: Socio-economic SCC-summary graph. Each node in this graph corresponds to a potentially persistent socio-economic profile. Node labels correspond to the socio-economic profiles. The “v” preceding a letter indicates the variation (i.e. +/- alternation) of the corresponding activity. An activity not mentioned is always absent of the set of states (e.g. in “AP”, T is absent). Node colors in each graph correspond to the same vegetation type. Arcs labels, exemplified by: R1|R3 means “Rule 1 or Rule 3”. Rule descriptions are detailed in Table S1.

#### Vegetation

The stable state corresponds to a desert-like ecosystem state (i.e. no vegetation, soil or wildlife). When starting in rainy season (Fig. S3), the HTG displayed a few large transient SCCs (≥ 100 states each), while most SCCs were small (<five states). Conversely, when starting in dry season, the smaller HTG (119 states, five SCCs) showed stable vegetation types in all SCCs (Fig. 3a,b). When projected onto a PCA plan, we observed that SCC profiles were discriminated along a land degradation and a grassland-savanna-woodland gradient, on the first and second axis, respectively (Fig. 3b). The irreversible dynamics mostly exhibit a trajectory along the first axis, from vegetated to bare soil.

#### Human activities

Agriculturalists communities persisted in all SCCs (Fig. 3a) before collapsing in the stable state. A socio-economic SCC-summary graph (Fig. 3c) allowed focusing on persistent socio-economic profiles in each vegetation type. Shifts in socio-economic profile were reversible within all vegetation type but bare soil.

### “Permanent high water availability” version (M2)

Removing the dry season rule induced, as in the M1 version, irreversible transitions (Fig. 4, S2 and S4). The HTG showed a large initial SCC in which several transitions in socio-economic profiles and vegetation could occur (blue node, Fig. 4), 294 transient SCCs (TS, Fig. 4 and Fig. S2) and two terminal SCCs (attractors A1 and A2). The two attractors were both reachable from the initial and transient SCCs. To highlight transitions leaving the initial SCC and those reaching the attractors, we compacted the 294 non-initial transient SCCs into one node (TS, Fig. 4), yielding a simplified HTG.

**Fig. 4.**
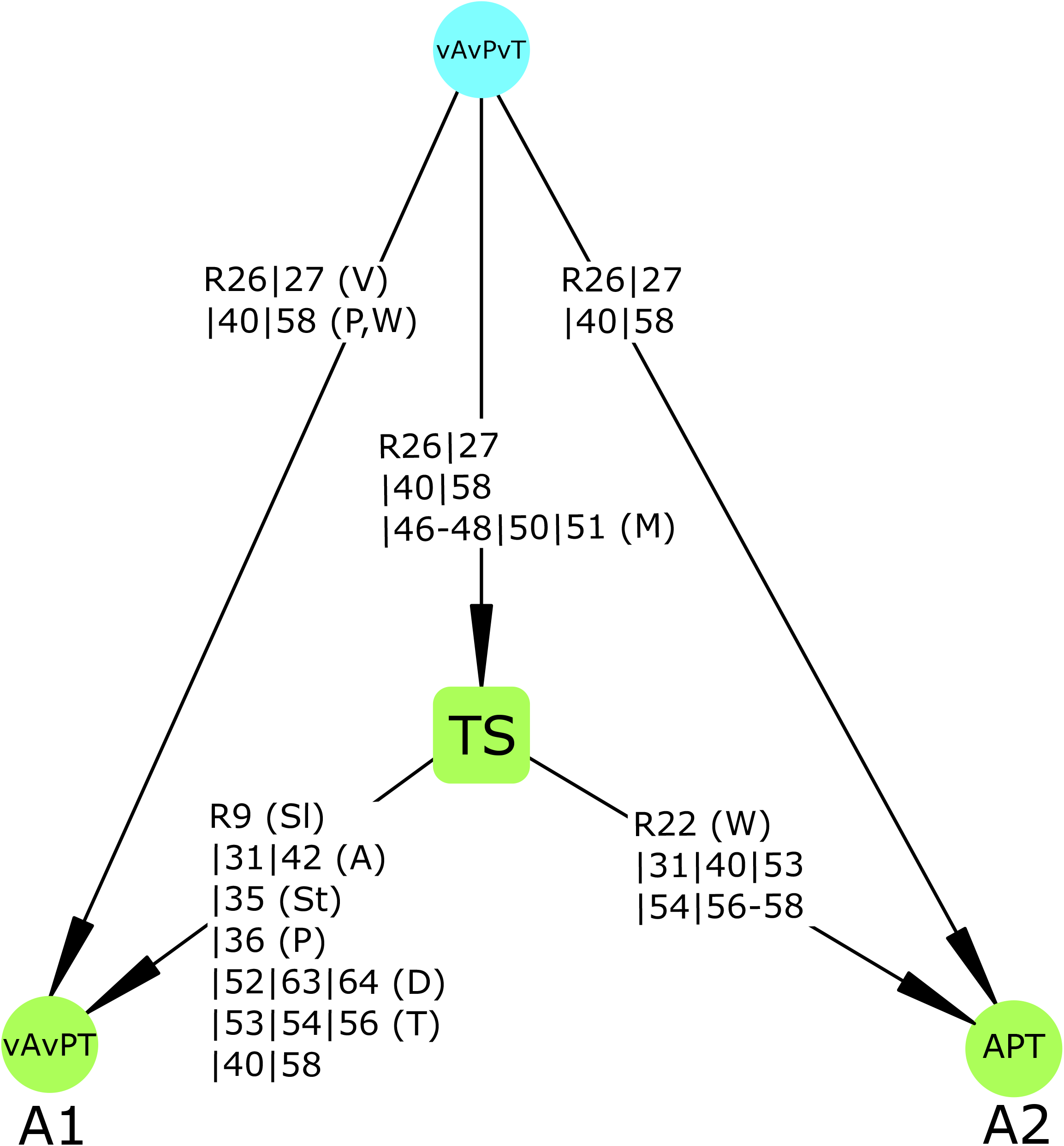
Sustained wet conditions (M2 model version). Simplified HTG, where each node is a set of states (SCC or basin). Round nodes are SCCs, among which A1 and A2 are attractors (terminal SCCs). The rounded square is an summary of all non-initial transient SCCs. Node colors represent vegetation dynamics: the blue node corresponds to a SCC including reversible grassland-savanna-woodland transitions; green nodes correspond to grassland only. Node labels correspond to socio-economic profiles, in which A corresponds to agriculture; P: pastoralism; T: tourism. The “v” preceding a letter indicates the variation (i.e. +/- alternation) of the corresponding activity in the set of states. Arcs labels, exemplified by: R1-3 means “Rule 1 or Rule 2 or Rule 3”; R1|R3 means “Rule 1 or Rule 3”. Each rule categorized based on its effect: V: vegetation; P: pastoralism; W: wildlife; M: management; Sl: soil; A: agriculture; St: settlements; D: diseases; T: Tourism. Rules categories are explained once, but rules may appear several times. Rules descriptions are detailed in Table S1.

#### Vegetation

Despite the high water availability, all SCCs but the initial one displayed a grassland vegetation type (green nodes) kept open by elephants/mixed-feeders (R26, Table S1 and S4). Strict browsers responded negatively to this increased water availability (not shown). The central role of elephants in the presence of irreversible transitions is supported by the reversibility of this model version when elephants are removed (not shown). In the latter case, the ecosystem may oscillate between different vegetation types. This scenario of delayed dry season was also assessed (Fig. S4).

#### Human activities

Attractors showed contrasted socio-economic dynamics. In A1, all activities were stable, whereas in A2, only tourism was stable while agriculture and pastoralism varied. These dynamics also share parts of their causal histories. Indeed, most transitions from TS to A2 were in common with TS to A1. Some TS-A1 transitions were related to soil formation (R9) or fight against wildlife and livestock diseases (R63-64), while most were related to socio-economy (e.g. R31, 53 or 40). Their last common direct predecessor SCC was the initial SCC, which indicates an early divergence. However, the two attractors share many indirect predecessors among the transient SCCs in TS, reflecting a non-deterministic “choice” process.

### “Grazers and browsers removal” versions (M3 and M4)

Compared to the reference model, removing grazers (M3) or browsers (M4) did not affect the reversibility of the STG. Initially induced by herbivory, plant mortality became induced by drought, thus preserving original vegetation transitions (Fig. 1c). Regarding socio-economy, major transitions related to pastoralism and agriculture were unchanged compared to reference model. Socio-economic transitions in M3 were the same as in Fig. 2d. However, in M4 (no grazers), tourism became impossible in absence of temporary water (Tw-). Indeed, as browsers are more drought resistant than grazers, the latter migrate earlier in dry season, thus reducing wildlife density and thus tourism. This in turn made transitions APT → AT and AT → NH (see Fig. 2d for labels) impossible. Consequently, as tourism now relies on browsers, their absence puts tourism at risk in case of moderate drought due to the absence of grazers (Tw-).

## Discussion

### The social-ecological system dynamics

Ecological models rarely address the structural dynamics of ecosystem. Rather, most models keep ecosystem structure and parameter values fixed. However, as (Jørgensen, 2015) puts it, “we know that an ecosystem can regulate, modify, and change [parameters], if needed as response to changes in the existing conditions”. The EDEN modelling approach presented in this paper is a tentative to account for such structural dynamics. Indeed, the discrete-event, qualitative savanna model presented here aimed to describe the possible qualitative states of a generic semi-arid and fireless East-African savanna. We identified all possible system states reachable under different model versions representing contrasting climate scenarios. Variables and rules were identified based on (i) direct field observations from two northern-Tanzanian savannas, (ii) expert knowledge and (Iii) literature about savannas across Africa (Devine et al., 2017; February et al., 2013; Homewood and Rodgers, 2004; Scholes and Archer, 1997; Sinclair, 2008).

We found that each social-ecological transition can ultimately be reversed (any state in the STG has the *possibility* to reach any other, i.e. it is a SCC) (Fig. 2a). As the model included a large number of variables and transitions, including different disturbances (epizootics, droughts, elephants browsing and livestock overgrazing), this result suggests that the overall ecosystem is *potentially* resilient. More precisely, after any disturbance, there exists at least one path leading to the pre-disturbance state. This definition of resilience does not take (1) recovery time, (2) system’s memory, (3) spatial heterogeneity, (4) the appearance of new variables nor (5) transitions probabilities into account. The potential resilience property is relatively independent of the initial state. Indeed, by definition of a SCC, if it holds for one initial state, then it holds also for any reachable state in this SCC. This implies that every vegetation shift is possibly reversible in this social-ecological system. This result is consistent with the literature on savannas dynamics pointing the existence of alternative stable states (grassland, woodland or savanna states; Baudena et al., 2015; Baxter and Getz, 2005; Briske et al., 2005), which can be reversed over long time scales (Alvarado et al., 2015; Gil-Romera et al., 2010; Sala and Maestre, 2014; Skarpe, 1992). Moreover, this reversibility can be extended to any aspect of the ecosystem (e.g. socio-economic profiles or livestock herds composition) in a context of normal rainy-dry seasonality (Fig. 2) (Skarpe, 1992, Sinclair, 2008).

### Vegetation dynamics

Among vegetation types, savanna had the largest social-ecological range (Fig. 2a and c, Table S6). This latter notion is comparable to that of Grinnellian ecological niche (Grinnell, 1917), for whom a niche is defined as the set of conditions of existence of a particular species. Although finding mostly savanna states may be congruent with their spatial prevalence in East-Africa, throughout Sub-Saharan Africa and other tropical and subtropical regions worldwide (Sankaran et al., 2005; Scholes and Archer, 1997), this model cannot confirm such a link due to its non-spatial and non-temporized nature.

The various model versions indicate that removing seasonality (versions M1, M2), herbivore communities (M3, M4) or anthropogenic variables (M5, M6, Supplementary Material), did not qualitatively modified vegetation ranges (Table S6). The fact that nomadic pastoralism does not strongly modify the vegetation (M6, Table S6) agrees with some findings (Homewood et al., 2009) but contrasts with others identifying it as a driver of bush encroachment (Sankaran et al., 2005). This low sensitivity can be explained by more transitions occurring in savanna. Indeed, wildlife and vegetation dynamics are much detailed parts of the model (i.e. driven by more transitions rules). We suggest that this greater detailed description leads to the observed combinatorial explosion in the number of savanna states. Another explanation could be that savanna states enable more herbivorous functional groups to coexist than less diverse states (i.e. grassland or woodland), thus resulting in more savanna. However, the choice of such herbivores functional groups and their diversity in mixed-vegetation was well supported by many studies (Cromsigt et al., 2009; Kartzinel et al., 2015). Nonetheless, assessing the role of pastoralism on vegetation in this model would require a more detailed analysis of trajectories.

### Socio-economic dynamics

Among socio-economic profiles, we found that agriculture-pastoralism combination has the largest social-ecological range (Fig. 3). As for savanna vegetation, this emergent observation has not been predefined in the model building and is in agreement with the literature since agriculture-pastoralism association started circa 5,000 years ago in East-Africa (Smith, 2005) and contributed to shaping African rangeland systems. As for the previous results on vegetation, this model cannot confirm such a link between results and observations due to its non-spatial and non-temporized nature.

Overall, pastoralism had the largest range of all human activities. This result fits well with the current knowledge about such tropical social-ecological systems, and especially for Maasailand rangelands, since pastoralism in these semi-arid lands is highly adapted to water fluctuations and vegetation types (Smith, 2005). All socio-economic profiles including tourism had less states than their no-tourism counterpart did (i.e. AP, P and A had more states than APT, PT and AT, respectively). This is probably because tourism is less detailed (i.e. involving fewer rules) than other activities such as pastoralism, leading to fewer states.

Permanently removing the rainy season (i.e. simulating a sustained drought scenario) induced the collapse of the system, which converged to a desert-like stable state (Fig. 3). We found drought and pastoralism to be the two primary proximal causes of desertification (here defined as any widespread environmental degradation that reduces productivity of dryland ecosystems by reducing plant cover and soil loss) (Peters et al., 2012; Schlesinger et al., 1990; Tucker et al., 1991). In this scenario, while tourism and pastoralism may vary, agriculture was stable and disappeared last (Fig. 3c). Indeed, the presence of bare soil did not preclude the presence of crop fields, which are primarily constrained by groundwater availability, as it can be used for irrigation (Amjath-Babu et al., 2016; de Bont et al., 2019). This results is not surprising, as pastoralists generally migrate towards wetter areas before the end of crop production (Agnew et al., 2001), in particular dry season agriculture which relies on water reserves in dry season (Nonga et al., 2011). Finally, this drought scenario contrasts with empirical results which suggest that desertification cannot be attributed to isolated factors, but rather to complex causal chains (Geist and Lambin, 2004). In comparison, the – more realistic - reference model, which provides many paths to desertification, may account for the multifactorial nature of this phenomenon. In this latter model, desertification was reversible (Fig. 2a, c) and the comparison with M1 confirms that rainfall strongly conditions this recovery (Peters et al., 2012).

Unexpectedly, removing the dry season (M2) was not a sufficient condition to maintain a woody or mixed vegetation in any case (Fig. 4). Although this is a direct consequence of the logical structure of the model, this was not intended during the modelling process and clearly emerged from the complex interplay between rules. Indeed, once the system exited the first large SCC, it was impossible for woody plants to regenerate. This irreversible vegetation transition was due to elephants and mixed-feeders, which became sedentary due to permanently sufficient water availability (Augustine and McNaughton, 2004) and prevented trees growth. This indicates that elephants, in this specific context, may determine the hysteresis of vegetation transitions. Such a control of tree regeneration by elephants is in agreement with previous studies (Baxter and Getz, 2005; Moe et al., 2009) and empirical observations as in the Mara Reserve in Tanzania since the 1970s (Dublin, 1986; Sinclair et al., 2008) or the Amboseli National Park in southern Kenya since the 1950s (Western and Maitumo, 2004). Unexpectedly, elephants seem to have contributed to a shift in socio-economic dynamics, by promoting either a highly diverse or a tourism-based economy. This suggests that in some conditions, more than just ecosystem engineers, elephants could be seen as “social-ecological engineers”. This observation points out the potential effect of biodiversity on the socio-economic dynamics, possibly through hardly predictable causal chains (TEEB, 2010). These results highlight the critical need for decision-makers to consider, not only anthropic or vegetation properties, but the whole network of biotic and abiotic variables in an integrative way to improve rangeland management through identification of their dominant stabilities and transitions (Borrelli et al., 2015).

Finally, removing grazers (M3) or browsers (M4) did not affect the general properties of vegetation dynamics, as transitions between bare soil, grassland, savanna and woodland were still possible. Indeed, in the reference model, vegetation changes were due to herbivory and drought (water), thus leaving drought as the sole driver in these two scenarios. While drought may induce major – but short-term – changes in grass biomass (Breshears et al., 2016; Wigley-Coetsee and Staver, 2020), a major reduction in trees biomass (Case et al., 2019) is likely (see Fensham et al., 2019 for reported historical evidence) and has been suggested as a possible mechanism for state transition to grass-dominated states (Breshears et al., 2016). Such transitions could profoundly and lastingly modify rangeland quality (Angassa, 2002). This leads scientists and managers in drought-prone areas to search for adaptive management strategies to cope with drought-related transitions (Bradford and Bell, 2017). On the socio-economic perspective, removing grazers did not affect major transitions and yielded the same results as in the reference model (Fig. 2d). As in the vegetation case, the number of processes responsible for socio-economic transitions was reduced. However, removing browsers affected tourism. As browsers have a greater resistance to drought (Abraham et al., 2019), their absence made tourism impossible in case of moderate drought (i.e. Tw-, once temporary water bodies have dried up) because grazers had already migrated. From a management perspective, this highlights the significance of maintaining small water bodies in dry season and a high mammal diversity for tourism, which may contribute to local incomes during the driest periods of the year where agriculture is mostly absent.

### An integrated qualitative model of a savanna social-ecological system

This study showed how discrete-event models allow for an integrated representation of ecosystem dynamics. Based on complex interaction networks and entangled causal chains, these models are able provide a synthetic view of ecosystem dynamics, promoting the dialogue between researchers and managers. This is also encouraging to understand ecosystem long-term dynamics and responses to disturbances. For instance, we found that maintaining a high water availability (M2, Fig. 4) could reduce ecological diversity by promoting a stable vegetation (Hirota et al., 2011) and herbivore communities (Fritz et al., 2002), while potentially increasing the socio-economic diversity. This highlights the potential of these models to assess the impact of planning procedures and the interplay between ecosystems and economics (TEEB, 2010). Besides the trajectories and terminal SCCs, the model provides the conditions under which a given variable or profile may be found and the trajectories in which it may persist or disappear. These tools are promising for identifying the transitions driving the stability of a complex system, allowing for an integrated sustainability assessment.

Our model provided many alternative system trajectories, which were disentangled using summary graphs, which rely on the aggregation of ecosystem states sharing some characteristics (e.g. vegetation or socio-economic profile). They thus allowed proposing answers about major ecosystem transitions, their causes and some long-term consequences. To allow a more detailed model analysis and following previous works in biology (Monteiro et al., 2008), oncoming developments focus on testing hypotheses and asking complex questions about trajectories using temporal logic (Clarke et al., 2018). Other improvements can be one in the chosen variables and rules. For instance, the model emphasized the role of water use in tree-grass interactions (Devine et al., 2017; February et al., 2013; Treydte et al., 2009). However, we could have modelled tree-grass interactions more finely by considering additional variables or processes such as fire or soil nutrients partitioning. These new processes could have generated new feedbacks in those interactions and modified the potential dynamics. Another limitation was the mixing of transitions of various spatio-temporal scales. However, ecologists have demonstrated the predominant role of spatial scales and heterogeneity in arid ecosystem dynamics (Klausmeier, 1999). Moreover, as untimed dynamics did not exhibit alternative SCCs (Fig. 1a), considering transitions durations may reveal different stability properties for each vegetation type (Skarpe, 1992). Thus, we could have instead characterized and ordinated transitions and variables in a spatially and temporally structured model. As a consequence, a spatially explicit extension of the model is currently developed (Leloup et al., In prep.). Regarding the temporal structure of the model, we are currently developing alternative methods to incorporate time constraints using temporal logic.

## Conclusion

With this study, we propose a discrete-event model predicting all the possible dynamics of a social-ecological system, represented as a States-Transitions Graph. The qualitative and possibilistic nature of the EDEN modelling approach accounts for generic and context-dependent interactions. It allows investigating causal pathways, which could support decision-makers when they are confronted to a high uncertainty. Moreover, EDEN has the potential to generate State-and-Transition Models automatically for the study and management of many ecosystems, including rangelands. We expect this approach to be promising for drawing a more dynamical view of social-ecological systems and supporting policy-makers in achieving both biodiversity conservation and food security persistence across Africa.

## Acknowledgements

We thank the HATARI project funded under the CNRS Tellus-Rift program for his financial support. We warmly thank Colin Thomas for his highly valuable comments and refinements of many ideas in the manuscript.

