## Supplementary Material for "East-African savanna dynamics: from a knowledge-based model to the possible futures of a social-ecological system"

**Appendix S1: Study area description**

Gelai plains are located in the East African Rift, south of Lake Natron, and are part of the Tarangire-Manyara ecosystem (Morrison and Bolger 2014) and the Maasailand. These grass-dominated plains have volcanic ash-derived alkaline soils. Indeed, their proximity with the Ol Doinyo Lengai stratovolcano, with its natrocarbonatite lava and ashes, strongly influences soil pH. Woody vegetation is confined to adjacent reliefs (Reed et al. 2009) and vegetation changes are mainly driven by water availability and herbivory (Sankaran et al. 2008; Asner et al. 2009). Mean annual precipitation strongly varies seasonally and inter-annually (Gereta et al. 2004). This high variability constrains primary, but also secondary production since the Gelai plains are important calving grounds for wildebeest populations (Morrison and Bolger 2014). The Gelai plains also provide resources for Maasai pastoralists and their cattle, goat and sheep herds. Agriculture is practiced but confined to village surroundings and wetter areas (Gereta et al. 2004). Lake Natron, mount Kitumbeine and Ol Doinyo Lengai are also important touristic sites, adding another economic value to this ecosystem.

On the other hand, Northern-Meru savannas are more densely populated. The different ethnics have contrasting economic activities, with characterized by the respective roles of agriculture and pastoralism during the year (Istituto Oikos 2011). Villages and local authorities (called land managers in the model) manage human activities by allowing water resources use, preventing overgrazing and allowing tourism. Their decisions also affect pathogen exchanges, vaccination, pest management and/or spatial and temporal grazing regulations. NGOs, such as Oikos, play a major role in improving livelihoods, promoting local development and sustainable practices. The area exchanges many goods and services with the Arusha city, south of the mount Meru.

**Appendix S2: Discrete-event modelling**

Transitions are an abstract representation of precedence, which is a qualitative property of time. Transition durations are implicit (and often unknown) and thus not represented. However, in this savanna model, transitions have a maximal extent (qualitative estimation) of about 100 years (e.g. soil formation) and a minimal resolution of approximately two weeks (e.g. shift from dry to early rainy season). Shorter processes are considered inevitable or irrelevant at the spatio-temporal scale considered, and thus represented as a constraint (i.e. a priority rule). Constraints generally represent transitions, which, at the ecosystem scale, result from strong interactions between ecosystem variables (e.g. a high reactivity of a variable to another one). They may represent a fraction of determinism in non-deterministic dynamics. Constraints do not have priority over each other.

For the abovementioned reasons, states satisfying a constraint are considered transient, and are therefore removed prior to analysis. The resulting state space is called the *compact state space*, which we simply referred to as *state space*. More details about state space computation have been published (Gaucherel and Pommereau 2019).

**Table S1. Rule set for the reference model version.** Variables abbreviations are provided in the main text (Table 2). "Field observations" may refer to direct or reported observations by local people, field experts or co-authors. "Common sense" rules refer to necessary relations regarding the variable's nature. Note that rule number differs between Reference model, M1 and M2 model versions (colored line).

| N° | Condition | Realization | Rule description | Reference |
| --- | --- | --- | --- | --- |
| R1 | Rf- | Rf+, Gr+, Tw+ | Rainy season | (February et al. 2013) and field observations |
| R2 | Rf+ | Rf- | Dry season | Common sense |
| R3 | Rf+ | Gr+, Tw+ | Frequent rainfall keeps grasses productive and surface water available. | Field observations |
| R4 | Rf+, Tw+ | Aq+ | In rainy season, aquifers may fill up (after surface water). | (Arnold et al. 1993) |
| R5 | Rf- | Tw- | In dry season, surface water may dry up. | Field observations |
| R6 | Rf-, Tw- | Aq- | In dry season, aquifers may dry up (after surface water). | (Arnold et al. 1993) |
| R7 | Rf-, Tw-, Lm+, Aq+ | Tw+ | In dry season, land managers may pump aquifers artificial waterholes from aquifers. | (Chamaillé-Jammes et al. 2016) |
| R8 | Rf-, Tw-, Lm+, Aq+ | Aq- | Filling artificial waterholes can lead to aquifers' draining. | (Kashimbiri et al. 2009) and assumption |
| R9 | Rf+, Gr+ | Sl+ | Moisture and grasses may promote soil formation. | (Maeda et al. 2010) |
| R10 | Rf-, Bv+, Lm- | Gr-, Sl- | In dry season, non-managed livestock may overgraze the area. | Schlesinger et al., 1990 |
| R11 | Rf-, Sh+, Lm- | Gr-, Sl- | In dry season, non-managed livestock may overgraze the area. | Schlesinger et al., 1990 |
| R12 | Ca+ | Gz- | Carnivores may strongly affect wild grazers populations. | Assumption |
| R13 | Ca+ | Bw- | Carnivores may strongly affect wild browsers populations. | Assumption |
| R14 | Ca+, Gz-, Bw-, St-, Lm- | Bv-, Sh-, Go- | Without managers support to protect livestock, carnivores may attack livestock if wildlife populations are low. | (Mponzi et al. 2014) |
| R15 | Lm-, Pa+ | Ca- | Without managers support to protect livestock, pastoralists may retaliate against carnivores. | (Mponzi et al. 2014) |
| R16 | Rf-, Tw- | El-, Bw-, Go- | Low water availability may lead browser and generalist herbivores to leave the area. | (Western 1975; Smit et al. 2007) |
| R17 | Bv+ | Gz- | Bovines may outcompete wild grazers. | Assumption |
| R18 | El+, Gr- | Go- | Elephants may outcompete goats. | Assumption |
| R19 | Go+, Gr- | El- | Goats may outcompete elephants. | Assumption |
| R20 | Go+ | Bw- | Goats may outcompete wild browsers. | Assumption |
| R21 | Bw+ | Go- | Wild browsers may outcompete goats. | Assumption |
| R22 | Gz+ | Ca+ | Wild grazers may increase carnivore populations. | Common sense |
| R23 | Bw+ | Ca+ | Wild browsers may increase carnivore populations. | Common sense |
| R24 | Tr+, Gr+, Rf+, Aq+ | Gr- | Trees outcompete grasses at high water availability. | (Devine et al. 2017) |
| R25 | Go+, Lm- | Tr- | Without grazing control, goats may strongly affect woody plants cover. | (O’Kane et al. 2011) |
| R26 | El+ | Tr- | Generalist herbivores may strongly affect woody plants cover. | (Laws 1970; O’Kane et al. 2011; Staver and Bond 2014) |
| R27 | Bw+ | Tr- | Browsers may strongly affect woody plants cover. | (Augustine and Mcnaughton 2004; Levick and Rogers 2008; O’Kane et al. 2011) |
| R28 | El+ | Bw- | Elephants may outcompete browsers. | (Fritz et al. 2002; O’Kane et al. 2011) |
| R29 | Rf+, Gr+, El-, Bw-, Go- | Tr+ | In rainy season, the absence of browsers may promote tree recruitment. | (Devine et al. 2017) |
| R30 | Rf-, Aq+, Tr+, Gr+ | Gr- | In dry season, trees maintain a higher productivity than grasses due to their deeper root system. | (February et al. 2013) |
| R31 | Rf+, Sl+ | Ag+, Cr+ | In rainy season, good soil conditions allow farmers to plant crops. | (Smaling and Dixon 2006; Maeda et al. 2010) |
| R32 | Rf-, Aq+, Lm+, Sl+ | Ag+, Cr+ | In dry season, land managers allow aquifers pumping for crop production. | (Kashimbiri et al. 2009) |
| R33 | Ag+, Tr+ | St+ | Farmers may build settlements from tree branches. | Field observations |
| R34 | Pa+, Tr+ | St+ | Pastoralists may build settlements from tree branches. | (Veblen 2013) and field observations |
| R35 | Pa-, Ag- | St- | Without humans, settlements may degrade. | Assumption |
| R36 | St- | Pa-, Ag- | Without settlements, humans may leave the area. | Assumption |
| R37 | Tw-, Aq+, Lm+, Tr+ | Go+, Pa+ | In dry season, land managers may pump groundwater for goats watering. | (Chamaillé-Jammes et al. 2016) |
| R38 | Tw+, Tr+ | Go+, Pa+, Bw+, El+ | Water and woody vegetation allow pastoralism and browsers establishment. | Field observations |
| R39 | Tw-, Aq+, Lm+, Gr+ | Bv+, Sh+, Pa+ | In dry season, land managers may pump groundwater for domestic grazers watering. | (Chamaillé-Jammes et al. 2016) |
| R40 | Tw+, Gr+ | Bv+, Sh+, Pa+, Gz+, El+ | Water and grasses allow pastoralism and grazers' establishment. | Field observations |
| R41 | Tr+ | Bw+ | Trees may promote browsers populations. | Field observations and (O’Kane et al. 2011) and common sense |
| R42 | Cr- | Ag- | Agriculturalists may leave if crops fail. | Assumption |
| R43 | El+, Lm- | Cr- | Without crop protection against elephants (e.g. fences), these may destroy crop fields. | Field observations and (Sinclair, 2008) |
| R44 | Ps+, Lm- | Cr- | Pests may destroy crops. | Field observations |
| R45 | Ag+, Lm- | El- | Without crop protection against elephants, farmers may kill elephants. | (Sinclair, 2008) |
| R46 | Pa+ | Lm+ | Pastoralism may induce/require management. | Assumption |
| R47 | Ag+ | Lm+ | Agriculture may induce/require management. | Assumption |
| R48 | Gz+ | Lm+ | Wildlife presence may induce management. | Assumption |
| R49 | Bw+ | Lm+ | Wildlife presence may induce management. | Assumption |
| R50 | El+ | Lm+ | Wildlife presence may induce management. | Assumption |
| R51 | Ca+ | Lm+ | Wildlife presence may induce management. | Assumption |
| R52 | Lm+ | Bd-, Gd- | Authorities may eradicate livestock and wildlife disease. | Field observations |
| R53 | Lm+, Gz+ | To+ | Wildlife and tourism monitoring may allow for tourism development. | Field observations |
| R54 | Lm+, El+ | To+ | Wildlife and tourism monitoring may allow for tourism development. | Field observations |
| R55 | Lm+, Bw+ | To+ | Wildlife and tourism monitoring may allow for tourism development. | Field observations |
| R56 | Lm+, Ca+ | To+ | Wildlife and tourism monitoring may allow for tourism development. | (Durant et al. 2011) and field observations |
| R57 | Lm+, Pa+ | To+ | Cultural activities (Maasai livelihood) and tourism monitoring may allow for tourism development. | Field observations |
| R58 | To+, Lm+, Gr+ | Ca+, Gz+, El+ | Tourism-derived income may contribute to wildlife conservation and restoration. | Field observations |
| R59 | To+, Lm+, Tr+ | Ca+, Bw+, El+ | Tourism-derived income may contribute to wildlife conservation and restoration. | Field observations |
| R60 | Rf+, Tw+, Go+, Bw+, Lm- | Bd+ | Without livestock/wildlife protection, high moisture and hosts contacts may promote browsers' and ovine pathogens. | (Hampson et al. 2011) |
| R61 | Rf+, Tw+, Sh+, Bw+, Lm- | Bd+ | Without livestock/wildlife protection, high moisture and hosts contacts may promote browsers' and ovine pathogens. | (Hampson et al. 2011) |
| R62 | Rf+, Tw+, Bv+, Gz+, Lm- | Gd+ | Without livestock/wildlife protection, high moisture and hosts contacts may promote browsers' pathogens. | (Hampson et al. 2011) |
| R63 | Bw-, Go-, Sh- | Bd- | Without hosts, pathogens may disappear. | (Hampson et al. 2011) |
| R64 | Gz-, Bv- | Gd- | Without hosts, pathogens may disappear. | (Hampson et al. 2011) |
| R65 | Lm+ | Ps- | Authorities may eradicate pests. |  |
| R66 | Rf+, Cr+ | Ps+ | In rainy season, pest may develop in crop fields. | (Gregory et al. 2009) |
| C1 | Bd+ | Bw-, Go-, Sh- | Browsers pathogens eradicate browser and ovine populations. | (Hampson et al. 2011) |
| C2 | Gd+ | Gz-, Bv- | Grazers pathogens eradicate grazer and cattle populations. | (Hampson et al. 2011) |
| C3 | Gz-, Bw-, Bv-, Sh-, Go- | Ca- | Without preys, carnivores disappear. | Common sense |
| C4 | Ca-, El-, Bw-, Gz-, Pa- | To- | Without cultural or wildlife attraction, tourists leave the area. | Common sense |
| C5 | Gr-, Tr- | El-, Bv-, Gz-, Sh-, Bw- | Without vegetation, herbivores disappear. | Common sense |
| C6 | Rf-, Tw-, Aq- | Ag-, Pa-, Bv-, Sh-, Gz-, Go-, Ca-, Bw-, El- | Without water, wildlife and humans disappear. | (Western 1975) |
| C7 | Rf-, Tw- | Pa-, Bv-, Sh-, Gz-, Ca- | Without surface water nor rainfall, grazers and pastoralism disappear. | (Western 1975) |
| C8 | Rf-, Aq- | Tr-, Gr-, Cr- | Crops and vegetation die during severe drought events. | Assumption |
| C9 | Pa- | Bv-, Sh-, Go- | Without pastoralists, livestock disappear. | Common sense |
| C10 | Gr- | Bv-, Gz-, Sh- | Without grass, grazers disappear. | Common sense |
| C11 | Tr- | Bw-, Go- | Browsers need woody plants. | Common sense |
| C12 | Pa-, Ag-, Bw-, El-, Gz- | Lm- | Without humans nor wildlife, management disappears. | Assumption |
| C13 | Gr-, Tr- | Sl- | Without vegetation, soil disappears. | (Durán Zuazo and Rodríguez Pleguezuelo 2008; Maeda et al. 2010) |
| C14 | Sl- | Cr- | Without soil, crops fail. | (Durán Zuazo and Rodríguez Pleguezuelo 2008) and field observations |
| C15 | Lm- | To- | Without management, tourism is not possible. | Assumption |
| C16 | Bv-, Sh-, Go- | Pa- | Pastoralists depend on livestock's survival. | Assumption |
| C17 | Ag- | Cr- | Without farmers, crops cannot persist. | Assumption |
| C18 | Cr- | Ps- | Without crops, pests cannot persist. | Assumption |

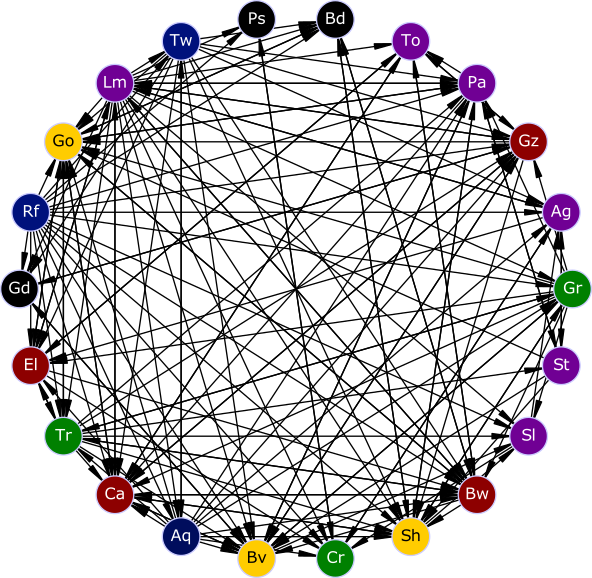

**Figure S1. Social-ecological interaction network.** An arc between two variables (nodes) indicates that the source variable is in the condition of a rule modifying the state of the target variable. Blue nodes: water-related variables; purple: humans and infrastructures; green: vegetation and crops; red: wildlife; black: pests and diseases; yellow: livestock; brown: soil. The disposition of nodes on the circle is arbitrary.

**Table S2. Transitions table of Fig. 2c (vegetation summary graph)**. Nodes correspond to vegetation types. Each arc (transition) is directed, and has a source (src) node and target (dst) node. A transition between two nodes may be caused by one or several underlying causes, resulting in several rules responsible for the transition considered. This implies that a STG, HTG or summary graph is formally defined as a directed multigraph. Rules descriptions are given in Table S1.

| src | dst | rules |
| --- | --- | --- |
| S | W | R30,R24,R11,R10 |
| S | G | R27,R25,R26 |
| S | B | R8,R6,R2 |
| W | S | R1,R3 |
| W | B | R27,R8,R26,R25,R6 |
| G | B | R8,R2,R11,R10,R6 |
| G | S | R29 |
| B | G | R1,R3 |

**Table S3. Transitions table of Fig. 2d (socio-economic summary graph)**. Nodes correspond to socio-economic profiles. Each arc is directed, and has a source (src) node and target (dst) node. A transition between two socio-economic profiles may be caused by one or several underlying causes, resulting in several rules responsible for the transition considered. This implies that a STG, HTG or summary graph is formally defined as a directed multigraph. Rules descriptions are given in Table S1.

| src | dst | rules |
| --- | --- | --- |
| APT | AT | R30,R27,R24,R21,R5,R26,R18 |
| APT | A | R30,R27,R24,R2,R5,R26 |
| APT | PT | R42 |
| APT | T | R36 |
| APT | NH | R36 |
| AT | NH | R8,R6 |
| AT | A | R30,R24,R5,R26,R27,R2,R16,R13,R12 |
| AT | APT | R38,R40 |
| AT | T | R36,R42 |
| A | NH | R36,R8,R5,R42,R6 |
| A | AP | R38,R40 |
| A | AT | R53,R56,R54,R55 |
| AP | APT | R53,R56,R54,R55,R57 |
| AP | A | R30,R60,R24,R21,R11,R5,R26,R10,R18,R27,R2,R14,R62,R61,R25 |
| AP | P | R42 |
| AP | NH | R36 |
| NH | P | R38,R40 |
| NH | A | R32,R31 |
| NH | T | R53,R56,R54,R55 |
| P | NH | R60,R24,R21,R11,R18,R36,R27,R62,R25, R30,R10,R5,R26,R2,R14,R61 |
| P | PT | R53,R56,R54,R55,R57 |
| P | AP | R32,R31 |
| PT | NH | R30,R36,R27,R24,R2,R5,R26 |
| PT | T | R30,R24,R21,R5,R26,R18,R36,R27 |
| PT | APT | R32,R31 |
| T | PT | R38,R40 |
| T | NH | R30,R24,R8,R5,R26,R27,R2,R16,R13,R6,R12 |
| T | AT | R32,R31 |

**Table S4. Model versions compared with the reference model**. Modifications are made on variables and/or rules. If a rule is removed in order to test the sensitivity of the model to this particular rule, this does not affect other rules or variables (e.g. M1 and M2). If a variable is removed, the rules involving this variable may be either (i) removed (in red) if they lose their ecological meaning (e.g. a rule in which elephants "*El*" destroy trees "*Tr*" makes no sense if elephants have been removed from the model) or (ii) modified according to the removed variable(s) (in blue). In this latter case, we only removed the removed variables from the rule if it stays an ecologically meaningful.

| **Model version** | **Modifications** | **Variables deleted** | **Rules deleted/modified** |
| --- | --- | --- | --- |
| M1 | Removing "Dry-to-rainy season" rule | None | R1 |
| M2 | Removing "Rainy-to-dry season" rule | None | R2 |
| M3 | Removing grazers variables and deletion/modification of associated rules | Wild grazers (Gz), Bovines (Bv), Sheeps (Sh) | R10-12, R14, R17, R22, R40, R48, R53, R58, R62, R63, R64, C3, C4-6, C8, C11, C13, C17 |
| M4 | Removing browsers variables and deletion/modification of associated rules | Wild browsers (Bw), Elephants (El) | R13, R16, R18-21, R23, R26-28, R29, R38, R40, R41, R43, R45, R49-50, R54-55, R58-61, R63, C1, C3-6, C11-12 |
| M5 | Removing anthropogenic variables and deletion/modification of associated rules | Pastoralists (Pa), Agriculturalists (Ag), Tourists (To), Land managers (Lm), St (Settlements), Crops (Cr), Bovines (Bv), Sheeps (Sh), Goats (Go) | R7-8, R10-11, R14-15, R16, R17-21, R25, R31-37, R38, R39, R40, R42-66, C1-3, C4, C5-8, C9, C10-11, C12-C18. |
| M6 | Removing pastoralism-related variables and deletion/modification of associated rules | Pastoralists (Pa), Bovines (Bv), Goats (Go), Sheeps (Sh) | R10-11, R14-15, R16, R17, R18-21, R25, R29, R34, R35-36, R37, R38, R39, R40, R46, R57, R60-64, C1-7, C9, C10-12, C16 |

**Table S5. M3 and M4 model versions rule sets.** Column "N°" indicates the rule (R) or constraint (C) number. Rules R1 to R9 have the same conditions and realization that the reference model, and are not detailed. Details about all rules (references and description) are given in Table S1.

| M3 | | | | M4 | | |
| --- | --- | --- | --- | --- | --- | --- |
| N° | Condition | | Realization | N° | Condition | Realization |
| R1 to R9 | | Identical to the reference model | | | | |
| R10 | Ca+ | | Bw- | Identical to the reference model | | |
| R11 | Ca+, Bw-, St-, Lm- | | Go- | Identical to the reference model | | |
| R12 | Lm-, Pa+ | | Ca- | Identical to the reference model | | |
| R13 | Rf-, Tw- | | El-, Bw-, Go- | **R13** | Ca+, Gz-, Bw-, St-, Lm- | Bv-, Sh-, Go- |
| R14 | El+, Gr- | | Go- | **R14** | Lm-, Pa+ | Ca- |
| R15 | Go+, Gr- | | El- | **R15** | Bv+ | Gz- |
| R16 | Go+ | | Bw- | **R16** | Gz+ | Ca+ |
| R17 | Bw+ | | Go- | **R17** | Tr+, Gr+, Rf+, Aq+ | Gr- |
| R18 | Bw+ | | Ca+ | **R18** | Rf+, Gr+ | Tr+ |
| R19 | Tr+, Gr+, Rf+, Aq+ | | Gr- | **R19** | Rf-, Aq+, Tr+, Gr+ | Gr- |
| R20 | Go+, Lm- | | Tr- | **R20** | Rf+, Sl+ | Ag+, Cr+ |
| R21 | El+ | | Tr- | **R21** | Rf-, Aq+, Lm+, Sl+ | Ag+, Cr+ |
| R22 | Bw+ | | Tr- | **R22** | Ag+, Tr+ | St+ |
| R23 | El+ | | Bw- | **R23** | Pa+, Tr+ | St+ |
| R24 | Rf+, Gr+, El-, Bw-, Go- | | Tr+ | **R24** | Pa-, Ag- | St- |
| R25 | Rf-, Aq+, Tr+, Gr+ | | Gr- | **R25** | St- | Pa-, Ag- |
| R26 | Rf+, Sl+ | | Ag+, Cr+ | **R26** | Tw-, Aq+, Lm+, Gr+ | Bv+, Sh+, Pa+ |
| R27 | Rf-, Aq+, Lm+, Sl+ | | Ag+, Cr+ | **R27** | Tw+, Gr+ | Bv+, Sh+, Pa+, Gz+ |
| R28 | Ag+, Tr+ | | St+ | **R28** | Cr- | Ag- |
| R29 | Pa+, Tr+ | | St+ | **R29** | Ps+, Lm- | Cr- |
| R30 | Pa-, Ag- | | St- | **R30** | Pa+ | Lm+ |
| R31 | St- | | Pa-, Ag- | **R31** | Ag+ | Lm+ |
| R32 | Tw-, Aq+, Lm+, Tr+ | | Go+, Pa+ | **R32** | Gz+ | Lm+ |
| R33 | Tw+, Tr+ | | Go+, Pa+, Bw+, El+ | **R33** | Ca+ | Lm+ |
| R34 | Tw+, Gr+ | | El+ | **R34** | Lm+ | Bd-, Gd- |
| R35 | Tr+ | | Bw+ | **R35** | Lm+, Gz+ | To+ |
| R36 | Cr- | | Ag- | **R36** | Lm+, Ca+ | To+ |
| R37 | El+, Lm- | | Cr- | **R37** | Lm+, Pa+ | To+ |
| R38 | Ps+, Lm- | | Cr- | **R38** | To+, Lm+, Gr+ | Ca+, Gz+ |
| R39 | Ag+, Lm- | | El- | **R39** | Rf+, Tw+, Sh+, Lm- | Bd+ |
| R40 | Pa+ | | Lm+ | **R40** | Rf+, Tw+, Bv+, Gz+, Lm- | Gd+ |
| R41 | Ag+ | | Lm+ | **R41** | Sh- | Bd- |
| R42 | Bw+ | | Lm+ | **R42** | Gz-, Bv- | Gd- |
| R43 | El+ | | Lm+ | **R43** | Lm+ | Ps- |
| R44 | Ca+ | | Lm+ | **R44** | Rf+, Cr+ | Ps+ |
| R45 | Lm+ | | Bd-, Gd- |  |  |  |
| R46 | Lm+, El+ | | To+ |  |  |  |
| R47 | Lm+, Bw+ | | To+ |  |  |  |
| R48 | Lm+, Ca+ | | To+ |  |  |  |
| R49 | Lm+, Pa+ | | To+ |  |  |  |
| R50 | To+, Lm+, Gr+ | | Ca+, El+ |  |  |  |
| R51 | To+, Lm+, Tr+ | | Ca+, Bw+, El+ |  |  |  |
| R52 | Rf+, Tw+, Go+, Bw+, Lm- | | Bd+ |  |  |  |
| R53 | Bw-, Go-, | | Bd- |  |  |  |
| R54 | Lm+ | | Ps- |  |  |  |
| R55 | Rf+, Cr+ | | Ps+ |  |  |  |
| C1 | Bd+ | | Bw-, Go- | **C1** | Bd+ | Sh- |
| C2 | Bw-, Go- | | Ca- | **C2** | Gd+ | Gz-, Bv- |
| C3 | Ca-, El-, Bw-, Pa- | | To- | **C3** | Gz-, Bv-, Sh- | Ca- |
| C4 | Gr-, Tr- | | El-, Bw- | **C4** | Ca-, Gz-, Pa- | To- |
| C5 | Rf-, Tw-, Aq- | | Ag-, Pa-, Go-, Ca-, Bw-, El- | **C5** | Gr-, Tr- | Bv-, Gz-, Sh- |
| C6 | Rf-, Tw- | | Pa-, Ca- | **C6** | Rf-, Tw-, Aq- | Ag-, Pa-, Bv-, Sh-, Gz-, Go-, Ca- |
| C7 | Rf-, Aq- | | Tr-, Gr-, Cr- | **C7** | Rf-, Tw- | Pa-, Bv-, Sh-, Gz-, Ca- |
| C8 | Pa- | | Go- | **C8** | Rf-, Aq- | Tr-, Gr-, Cr- |
| C9 | Tr- | | Bw-, Go- | **C9** | Pa- | Bv-, Sh- |
| C10 | Pa-, Ag-, Bw-, El- | | Lm- | **C10** | Gr- | Bv-, Gz-, Sh- |
| C11 | Gr-, Tr- | | Sl- | **C11** | Pa-, Ag-, Gz- | Lm- |
| C12 | Sl- | | Cr- | **C12** | Gr-, Tr- | Sl- |
| C13 | Lm- | | To- | **C13** | Sl- | Cr- |
| C14 | Go- | | Pa- | **C14** | Lm- | To- |
| C15 | Ag- | | Cr- | **C15** | Bv-, Sh- | Pa- |
| C16 | Cr- | | Ps- | **C16** | Ag- | Cr- |
|  |  | |  | **C17** | Cr- | Ps- |

**Versions with different sets of variables (M3-M6)**

As in the reference model, savanna and bare soil had the largest and smallest range of all vegetation types for all model versions, respectively (Table S6). Removing grazers or browsers increased the proportion of grassland states. Removing humans or pastoralism had the same effect on ranges but not on state space size, suggesting a reduced effect of pastoralism on vegetation dynamics. However, as mentioned in the main text, these results should be taken cautiously due to the contrasting combinatorial potential of the various variables.

**Table S6.** Quantification (approximate % of state space size) of the social-ecological range of conditions of existence (hereafter called range) of each vegetation type (G, S, W, and B) in the different versions of the model (M1-M6). G: grassland; S: savanna; W: woodland; B: bare soil. Parentheses indicate the removed rules or variables. Attractors refers to the presence and nature. s.s: single stability. Bold percentages indicate maxima.

| **Model version** | **G** | **S** | **W** | **B** | **Reversible state space** | **# states** |
| --- | --- | --- | --- | --- | --- | --- |
| Reference | 21% | **68%** | 10% | 0,4% | Yes | 24 444 |
| M1 | 20% | **67%** | 13% | 0,3% | No (one bare soil fixed state) | 119 |
| M2 | 13% | **52%** | 29% | 7% | No (two grassland attractors) | 8180 |
| M3 | 35% | **51%** | 12% | 1% | Yes | 3588 |
| M4 | 45% | **48%** | 5% | 1% | Yes | 4980 |
| M5 | 25% | **44%** | 25% | 6% | Yes | 352 |
| M6 | 27% | **50%** | 22% | 2% | Yes | 6456 |

**Table S7.** Socio-ecological range of each socio-economic profile in each vegetation type for the reference model version. Bold percentages indicate maxima.

| **Vegetation type** | **AP** | **APT** | **P** | **A** | **PT** | **AT** | **None** | **T** | **# States** |
| --- | --- | --- | --- | --- | --- | --- | --- | --- | --- |
| All | **33,1%** | 17,1% | 16,5% | 12,1% | 8,5% | 4,6% | 5,6% | 2,3% | 24 444 |
| Savanna | **36,2%** | 18,7% | 18,1% | 8,3% | 9,3% | 3,5% | 4% | 1,8% | 16712 |
| Grassland | **30,2%** | 15,6% | 15,1% | 15,7% | 7,8% | 5,6% | 7,1% | 2,9% | 5084 |
| Woodland | 26,9% | 13,4% | 13,4% | **37%** | 6,7% | 13,4% | 16% | 6,7% | 2544 |
| Bare soil | 0% | 0% | 0% | **62%** | 0% | 0% | 38% | 0% | 104 |

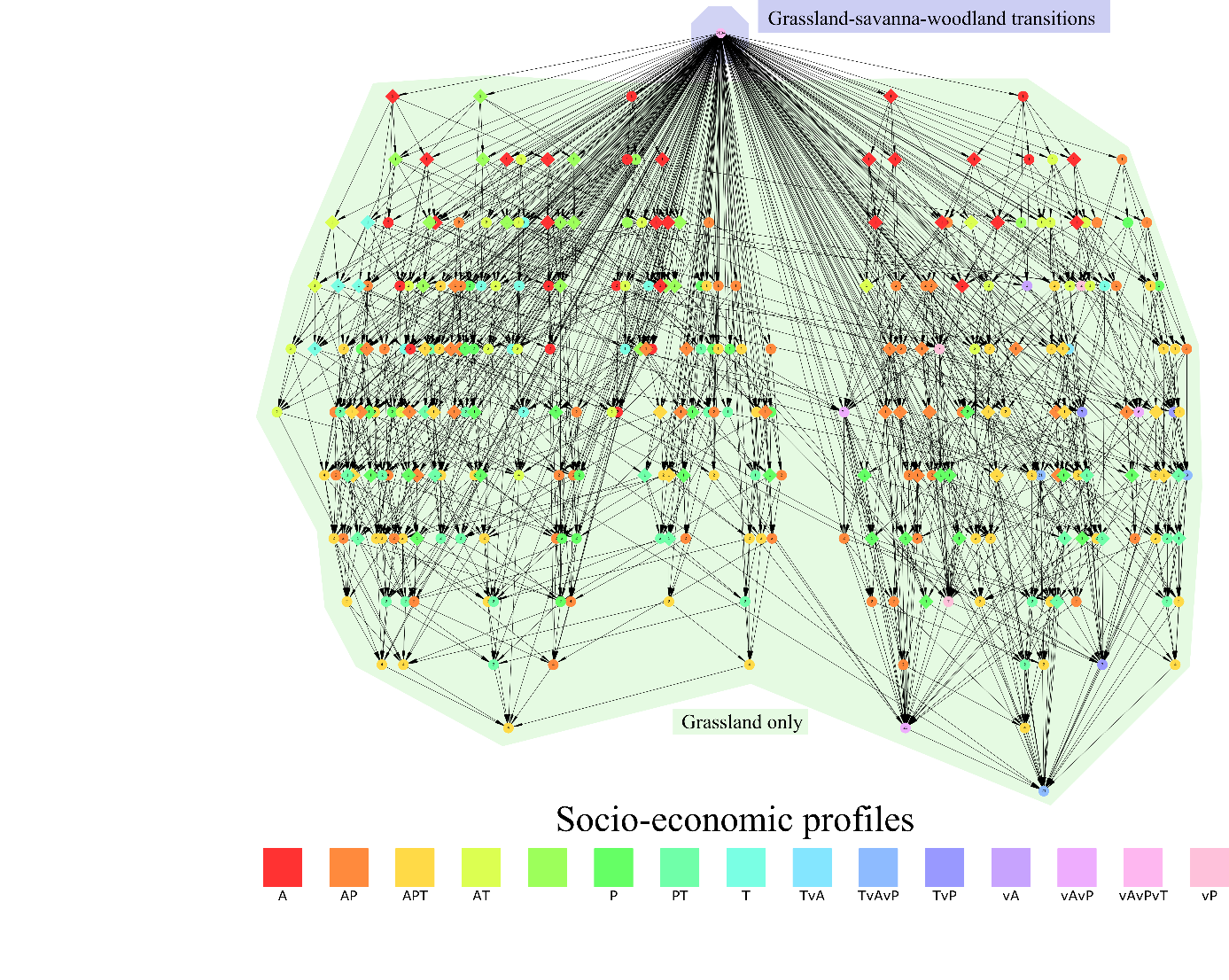

**Fig. S2**. **Permanent rainy season (M2 model version)**. Full merged state space. The initial state is in rainy season. Terminal stabilities are attractors. Blue and green shades correspond to varying (grassland, savanna or woodland) and grassland vegetation types, respectively. Node labels represent the size (in number of states) for each stability. Node colors correspond to socio-economic profiles, where A corresponds to agriculture; P: pastoralism; T: tourism. The "v" preceding a letter indicates the variation (i.e. +/- alternation) of the corresponding activity in the set of states.

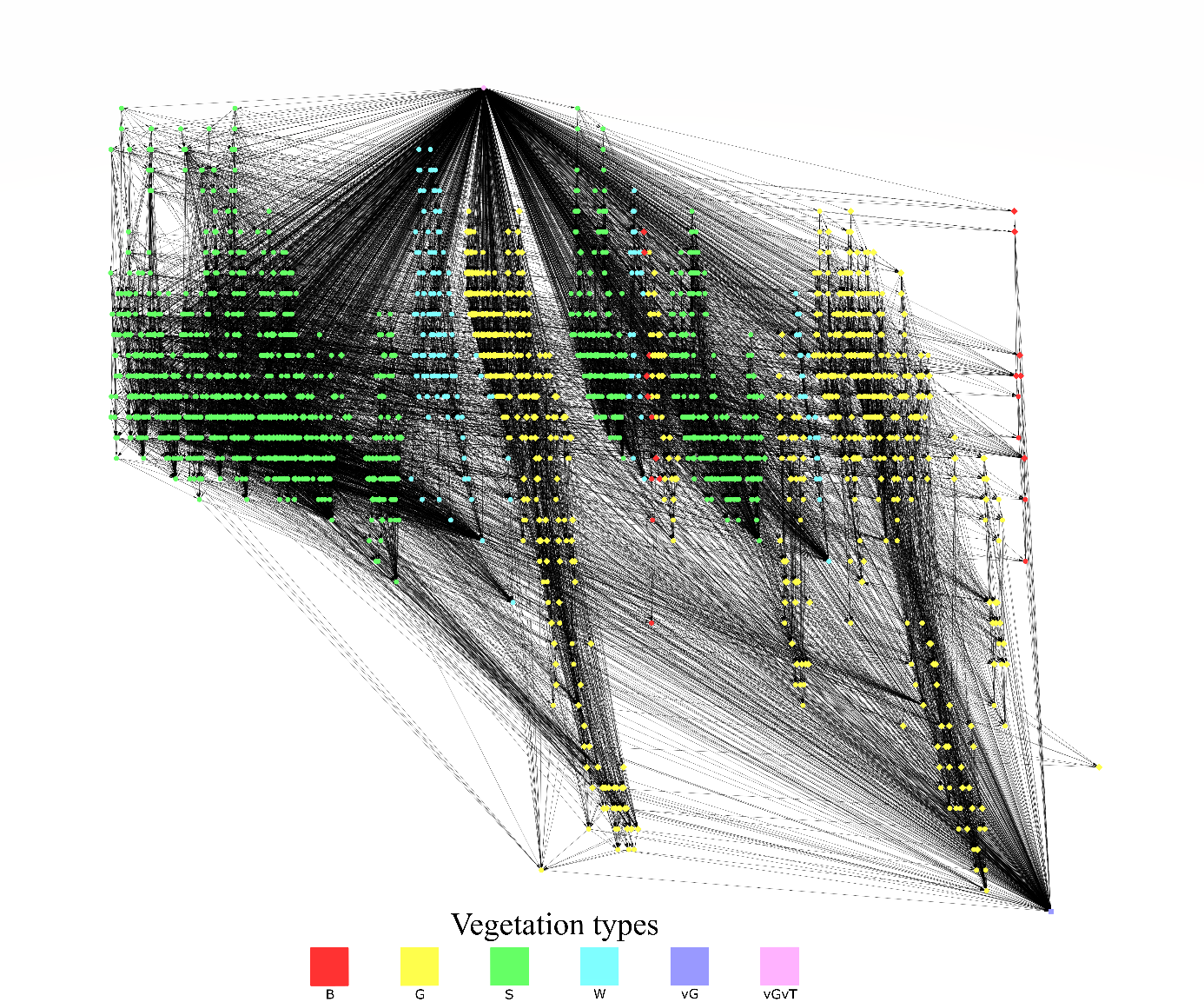

**Figure S3. Merged state space (M2 model version)**.The initial state is in rainy season. As in the main text, the attractor is a deadlock (fixed-state). Each node is a stability. Node colors refer to vegetation types. B: bare soil; G: grassland; S: savanna; W: woodland; vG: grassland with temporary bare soil; vGvT : alternations of the previously mentioned vegetation types. The "v" preceding a letter indicates the variation (i.e. +/- alternation) of the corresponding vegetation types in the stability.

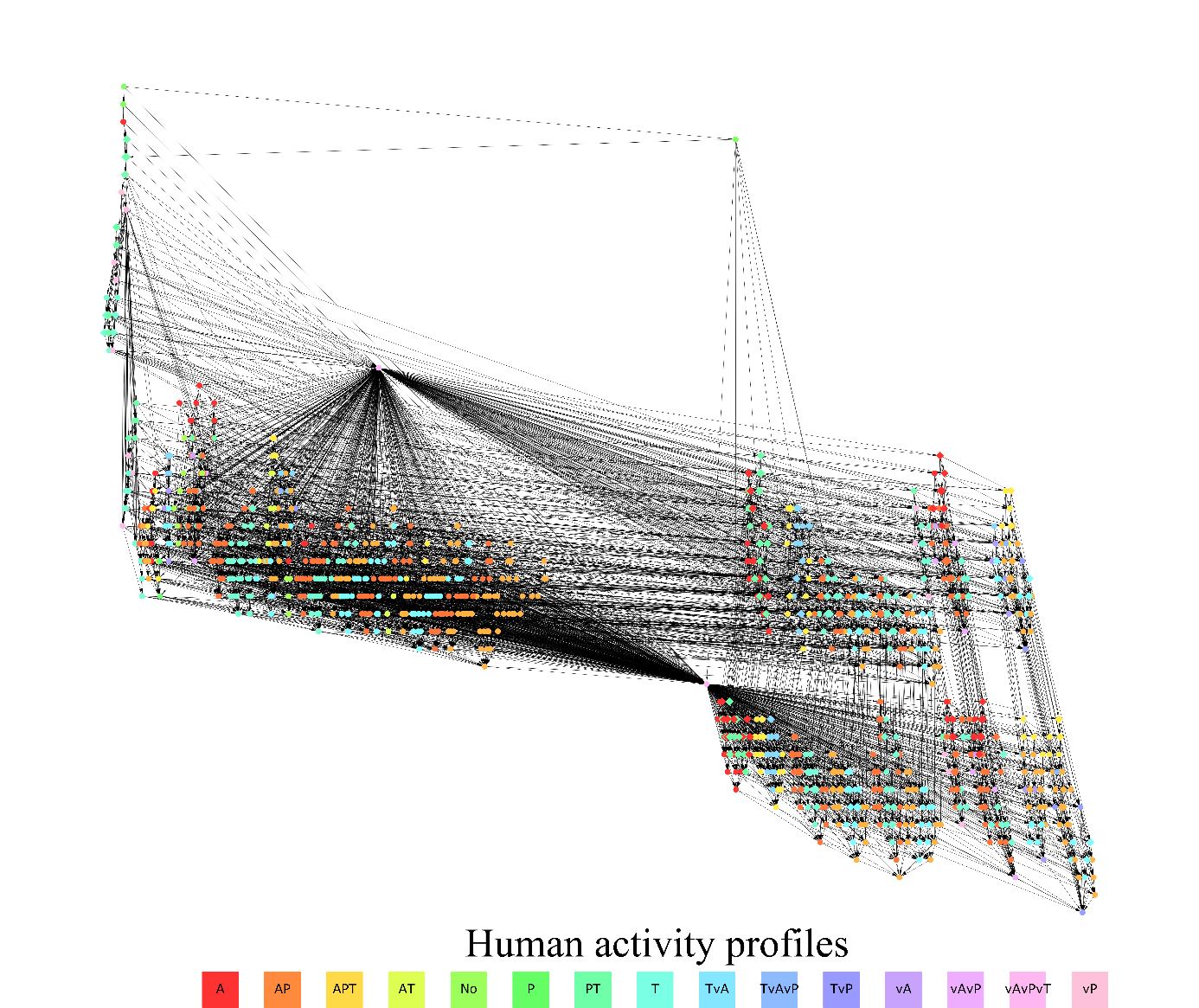
**Figure S4. Merged state space ( M3 model version).** The initial state is in dry season. The node colors correspond to activity profiles, made of (possibly varying) stable combinations of pastoralism (P), agriculture (A) and tourism (T). The "v" preceding a letter indicates the variation (i.e. +/- alternation) of the corresponding vegetation types in the stability. “No” corresponds to human absence. The two attractors are the same as in Fig. 4 in the main text.
